# Convergence within divergence: insights of wheat adaptation from *Triticum* population sequencing

**DOI:** 10.1101/2020.03.21.001362

**Authors:** Yao Zhou, Xuebo Zhao, Yiwen Li, Jun Xu, Aoyue Bi, Lipeng Kang, Haofeng Chen, Ying Wang, Yuan-ge Wang, Sanyang Liu, Chengzhi Jiao, Hongfeng Lu, Jing Wang, Changbin Yin, Yuling Jiao, Fei Lu

## Abstract

Bread wheat expanded its habitats from a small core area of the Fertile Crescent to global environments within ∼10,000 years. Genetic mechanisms of this remarkable evolutionary success are not well understood. By whole-genome sequencing of populations from 25 subspecies within genera *Triticum* and *Aegilops*, we identified composite introgression from these wild populations contributing 13%∼36% of the bread wheat genome, which tremendously increased the genetic diversity of bread wheat and allowed its divergent adaptation. Meanwhile, convergent adaption to human selection showed 2- to 16-fold enrichment relative to random expectation in *Triticum* species despite their drastic differences in ploidy levels and growing zones, indicating the vital importance of adaptive constraints in the success of bread wheat. These results showed the genetic necessities of wheat as a global crop and provided new perspectives on leveraging adaptation success across species for crop improvement.

## Introduction

Bread wheat (*Triticum aestivum*. ssp. *aestivum*) is one of the most successful crops on earth since the Neolithic Age. Within only a few millennia, wheat expanded its habitat from a small core area within the Fertile Crescent to a broad spectrum of diverse environments around the globe, making it the most widely grown crop in the world^1,2^. Despite, wheat serving as one of the keystone crops for global food security, offering ∼20% of calories and proteins of human diet^3^, new environments resulting from climate change may compromise its production^4,5^. Understanding the genetic mechanisms of the adaptation success is key to productive and stable wheat production in the future.

Bread wheat (2n = 6x = 42, AABBDD) is an allohexaploid species, comprising of A, B, and D subgenomes. It originated from two successive rounds of polyploidization within genera *Triticum* and *Aegilops*, forming tetraploid wheat (AABB) and hexaploid wheat (AABBDD), respectively^6^. By fusing genomes previously existing in different environments, polyploidization brought broader adaptability to bread wheat^7^. However, the same polyploidization process combined with the domestication bottleneck induced severe diversity reduction to the ancestral population of bread wheat^8,9^, which was likely impeding its range expansion. Recent studies identified that introgression from wild emmer (*T. turgidum*. ssp. *dicoccoides*, AABB) could increase the diversity of bread wheat^10,11^. Nevertheless, given the complex taxonomic groups within genera *Triticum* and *Aegilops* plus their unresolved phylogeny (Supplementary Table 1)^12-14^, understanding the diversity recovery of bread wheat from introgression may be still far from enough. In addition to the obscurity of the divergent adaptation, questions on the convergent side of the adaptation coin remain to be addressed. The evolution of wheats, or *Triticum* species, is deeply rooted in human selection for agronomic traits, which almost exclusively centered around food production. During the process, wild einkorn (*T. monococcum*. ssp. *aegilopoides*, AA) and wild emmer were independently domesticated. Likewise, durum wheat (*T. turgidum*. ssp. *durum*, AABB) and bread wheat were separately improved^12,13,15^. It is natural to ask how similar the wheat genomes were transformed to meet similar human needs during these selection events. Decoding the paralleled adaptation process across species, ploidy levels, and growing zones is crucial to identify the genetic necessities of bread wheat to produce large amounts of carbohydrates while growing in different environments, which is particularly valuable for wheat breeding during in a changing climate.

Here we report the first genus-level whole-genome sequencing study of wheat, in which all possible genetic donors of bread wheat within the genus *Triticum* were analyzed. Several subspecies from *Aegilops tauschii* were also sampled to study the evolution of D subgenome of bread wheat. The analysis provides novel insights about genetic driving forces behind wheat adaption with a highlight on evolutionary constraints of wheat adaptation to human selection. New perspectives from the study should be beneficial to wheat genetic research and breeding in the future.

## Results

### The worldwide collection of wheats from the genus *Triticum*

We sampled all species of AA, AABB and AABBDD genomes within *Triticum*, and the progenitor of D subgenome of bread wheat, *Ae. tauschii*. This collection consisted of *Triticum* species from a wide range of taxonomic groups, geographic distribution, and ploidy levels. A total of 414 accessions of 5 species, 25 subspecies from 71 countries were collected (Fig. 1a, Supplementary Table 2, for convenience purposes, common names of these subspecies were used in this study), including 91 diploid *Triticum* accessions (AA taxa), 30 diploid *Ae. tauschii* accessions (DD taxa), 121 tetraploid accessions (AABB taxa), and 172 hexaploid accessions (AABBDD taxa). In order to dissect paralleled selection events of domestication and improvement, paired sampling of einkorn and emmer (Fig. 1b), as well as durum and bread wheat (Fig. 1c) were elaborately conducted.

**Fig. 1.**
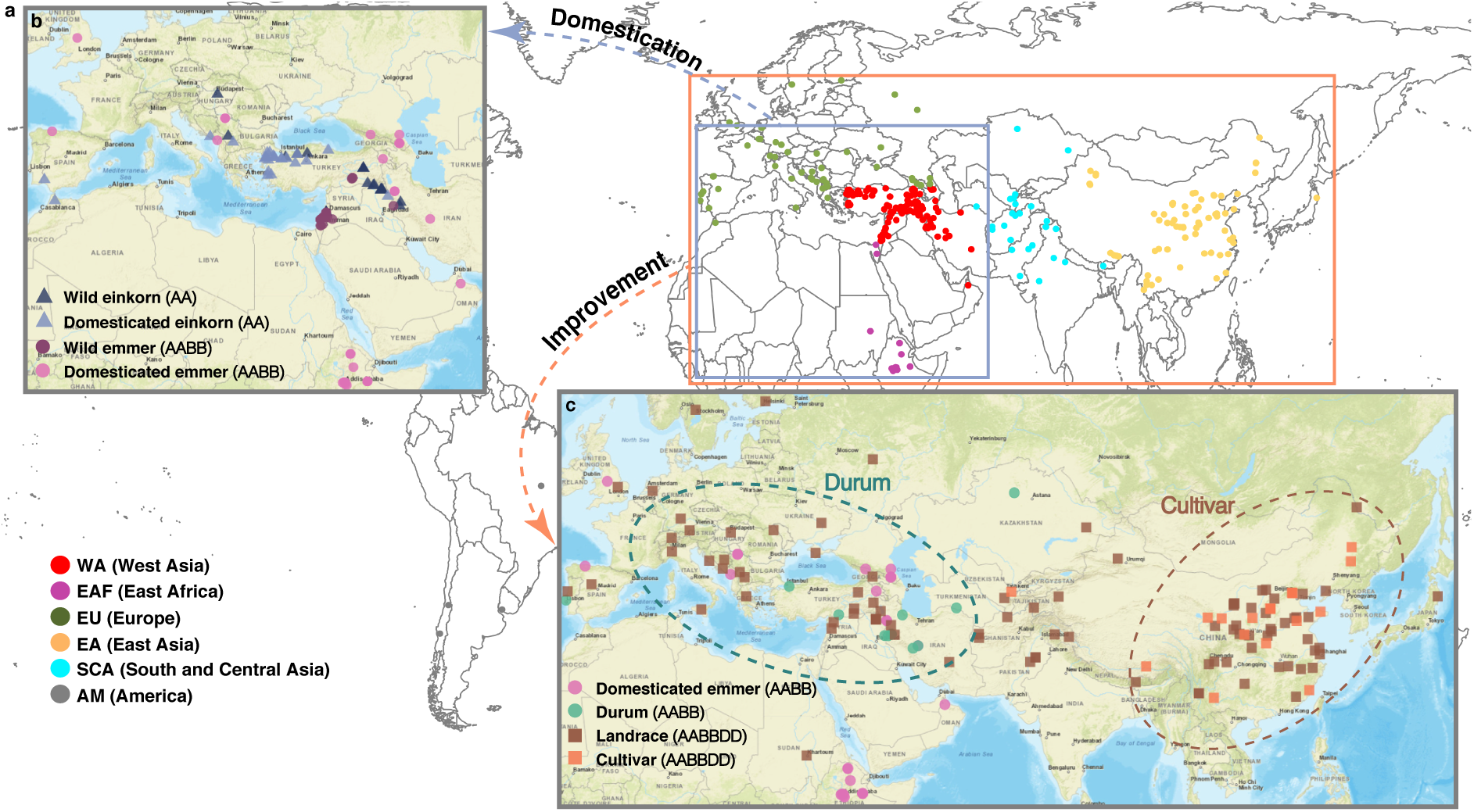
The worldwide collection of wheat accessions. **a**, Geographic distribution of all accessions. Dots with red, purple, green, brown, cyan, grey colors represent accessions from West Asia (EA), East Africa (EAF), Europe (EU), East Asia (EA), South and Central Asia (SCA), and America (AM). **b**, Accessions used for selection scan of wheat domestication events. Wild einkorn, domesticated einkorn, wild emmer, and domesticated emmer are color-coded. **c**, Accessions used for selection scan of wheat improvement events. Domesticated emmer, durum, landrace and cultivar of bread wheat are color-coded.

### A comprehensive whole-genome genetic variation map of wheat

To obtain genome-wide genetic variations and fully present adaptive signatures of bread wheat, we constructed a comprehensive whole-genome genetic variation map of wheat (VMap I) by whole-genome sequencing of all 414 accessions with an average coverage at 6x (Supplementary Table 3). A custom-designed pipeline for cross-ploidy variation discovery was implemented to overcome the multiple ploidy levels within the collection and ensure high accuracy of variation calling (Supplementary Fig. 1). First, genetic variations were discovered from each genome/subgenome group, e.g. AA taxa, by taking corresponding genome/subgenomes from the wheat genome IWGSC RefSeq v1.0^16^ as its own reference. Next, to avoid reference bias of cross-species variation calling, only genetic variations in syntenic regions within each lineage were retained (Supplementary Fig. 2a,b). Then, variations from each group were merged into VMap I. Respectively, about 13%, 44%, and 48% of genomic sites were defined as syntenic in A, B and D subgenome, which helped to identify a total of ∼104M single nucleotide polymorphisms (SNPs). In the final data set, the A, B, and D subgenomes contained ∼30M, ∼49M, and ∼25M SNPs, respectively. The error rate of variant calling, i.e., the proportion of segregating sites in the reference accession, Chinese Spring, was 0.024%, which was similar to the error rate of SNP calling in cassava^17^. The low error rate was also verified by the lowest genetic divergence (0.0037) between the sample of Chinese Spring and the reference genome compared with all other genotyped accessions (Supplementary Fig. 2c).

### Phylogeny of *Triticum* species

Phylogeny is fundamental to infer population structure and investigate adaptive changes of a species. However, lacking an accurate phylogeny of *Triticum*^12-14^ impeded active investigation on wheat adaptation. Therefore, with hundreds of thousands of randomly chosen SNPs, we applied a stepwise approach to obtain robust phylogenies of *Triticum* within A lineage (AA, AABB, and AABBDD taxa), AB lineage (AABB and AABBDD taxa), and D lineage (AABBDD and DD taxa).

With barley (*Hordeum vulgare*) as the outgroup, the phylogeny of the A lineage based on 200K SNPs showed that wild einkorn, domesticated einkorn, and urartu were explicitly separated from AABB and AABBDD taxa (Fig.2a and Supplementary Fig. 3a). Model-based estimation of ancestry from sNMF^18^ also demonstrated unique genetic composition of these taxa (Supplementary Fig. 3b). However, A subgenome specific SNPs represented merely half of the genome of AB lineage, and only 39% of them were segregating within AB lineage, due to the long divergence time of 0.82 million years between AA and AABB taxa^6^. To reconstruct evolutionary relationship of *Triticum* species within the most recent ∼10,000 years, we resampled 200K standing variations in AB lineage to obtain a set of unbiased and informative SNPs.

As indicated by the phylogeny of A lineage (Supplementary Fig. 3a), wild emmer accessions were set to the root of the phylogeny of AB lineage. AB lineage exhibited a continuum of ploidy levels, from tetraploids, then a mixture of tetraploids and hexaploids, to pure hexaploids, reflecting the tetraploid ancestry of bread wheat. The phylogeny of the AB lineage also presented a clear shift from hulled wheats to free-threshing wheats, indicating the vital role of human selection during the evolution of *Triticum* (Fig. 2b and Supplementary Fig. 4a,b).

**Fig. 2.**
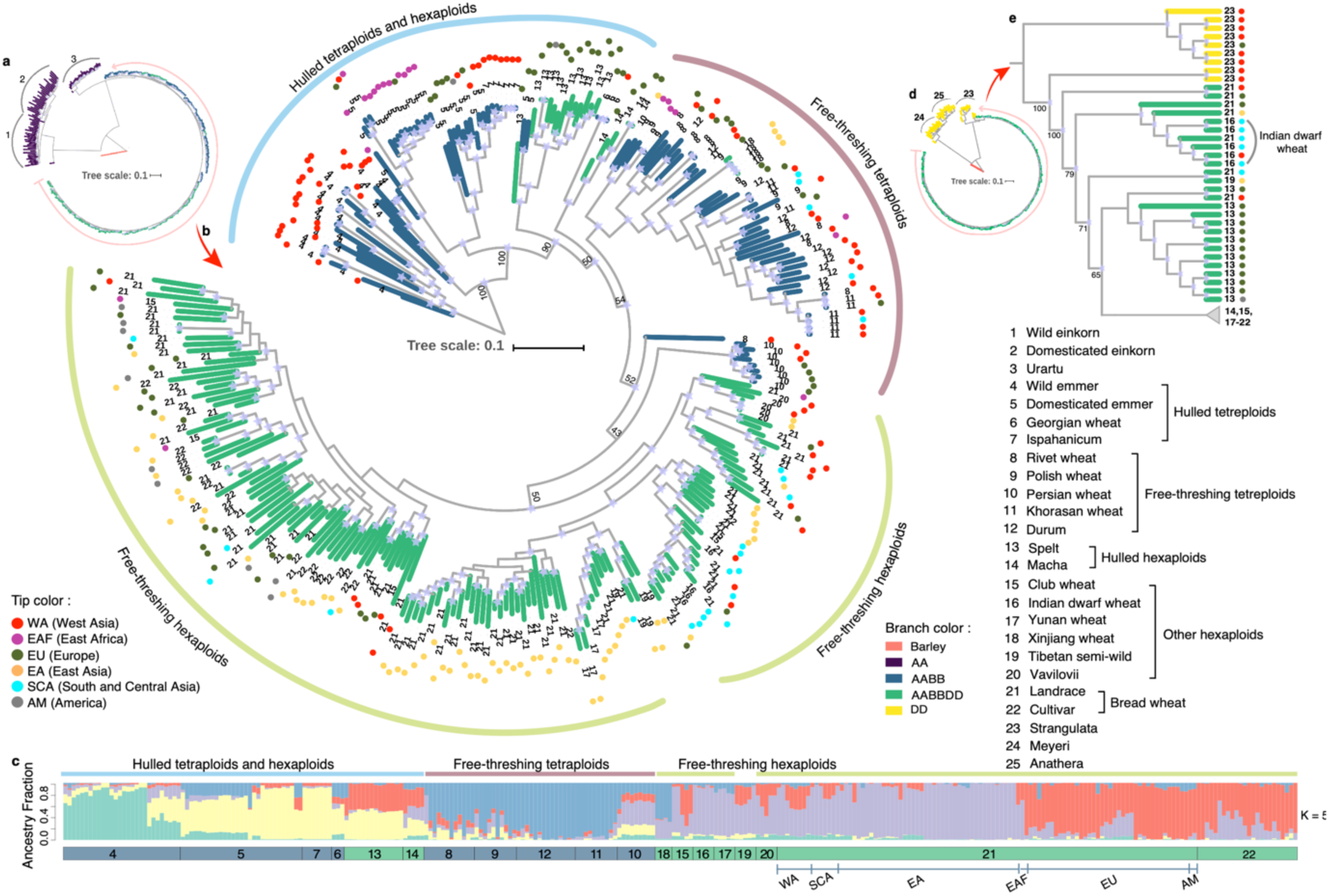
Phylogenetic relationship of *Triticum* species. **a**, Phylogeny of A lineage with barley as the outgroup. **b**, Phylogeny of AB lineages with wild emmer as the outgroup. **c**, Ancestry coefficient analysis of AB lineages with K = 5. **d**, Phylogeny of D lineage with barley as the outgroup. Clades with bootstrap values less than 50 were collapsed. **e**, Phylogeny of D lineage with strangulata as the outgroup. Clades with bootstrap values less than 50 were collapsed. For subfigures a, b, d and e, tip colors of the phylogeny are sampling regions of each accession, while branch colors are ploidy levels. Branches with reliable bootstrap values (> 50) were labeled with a purple pentagram at corresponding nodes. Numbers represent species/subspecies from 1 to 25.

The rise of bread wheat has long been controversial. There is a consensus that bread wheat and other free-threshing hexaploids originated from hybridization between tetraploid wheats and *Ae. Tauschii*. However, based on competing evolutionary models, the tetraploid donor could either be domesticated emmer^19^, free-threshing tetraploids^20^, or durum^21,22^. Our results showed that free-threshing tetraploids were the closest basal group to all bread wheat, including both bread wheat landraces (hereinafter referred to as landrace) and bread wheat cultivated varieties (hereinafter referred to as cultivar), indicating that the direct donor of AB subgenomes of bread wheat was mostly likely to be free-threshing tetraploids (Fig. 2b).

Landraces were clearly divided into Asian and European groups (Fig. 2b,c). Instead of being clustered with East Asian (EA) landraces, nearly 91% of East Asian cultivars (*n* = 22), mostly collected in China, were grouped with European landraces (Fig. 2b and Supplementary Fig. 4a). In line with this, both ancestry coefficient estimation and population differentiation statistics (*F*_*ST*_) showed that EA cultivars were genetically more similar to European landraces (Fig. 2c and Supplementary Fig. 4b,c,d). These results are consistent with a recent analysis showing that modern Chinese cultivars derive from the intensive use of European wheat^23^.

The phylogeny of D lineage confirmed that strangulata (*Ae. tauschii*. ssp. *strangulata*) was the D subgenome progenitor of bread wheat (Fig. 2d,e)^24^. In addition, we found that Indian dwarf wheat (*T. aestivum*. ssp. *sphaerococcum*), an endemic taxon to southern Pakistan and northwestern India, was basal to 96% of bread wheat in the phylogeny of D lineage (Fig. 2e and Supplementary Fig. 5). The basal phylogenetic position across hexaploids made Indian dwarf wheat a highly valuable reference population to investigate the recent adaptation of bread wheat.

### Genetic diversity of *Triticum* species

A comprehensive survey was conducted to evaluate nucleotide diversity (*π*) of bread wheat and its wild relatives (Fig. 3a and Supplementary Table 4), and further quantify the diversity reduction of bread wheat resulting from polyploidization and domestication^8,9^. The results showed that bread wheat had intermediate genome-wide nucleotide diversity (*π*) among major crops (*π*A = 0.0017, *π*B = 0.0025, *π*D = 0.0002), which was much lower than maize (*Zea mays*) (*π* = 0.014)^25^, but similar to rice (*Oryza sativa*) (*π* = 0.0024)^26^. Wild emmer maintained the highest nucleotide diversity across all taxonomic groups (*π*_A_ = 0.0025, *π*_B_ = 0.0042), together with its similar physiology with bread wheat, making it an invaluable gene pool for genetic improvement of wheat. Indeed, significant reductions in genetic diversity resulting from bottleneck events were observed. For example, domesticated emmer had 73% (one-tailed *t*-test, *P* = 3.4 × 10^−208^) of nucleotide diversity from wild emmer, while domesticated einkorn captured only 56% (one-tailed *t*-test, *P* = 8.4 × 10^−136^) from its wild form (Fig. 3a).

**Fig. 3.**
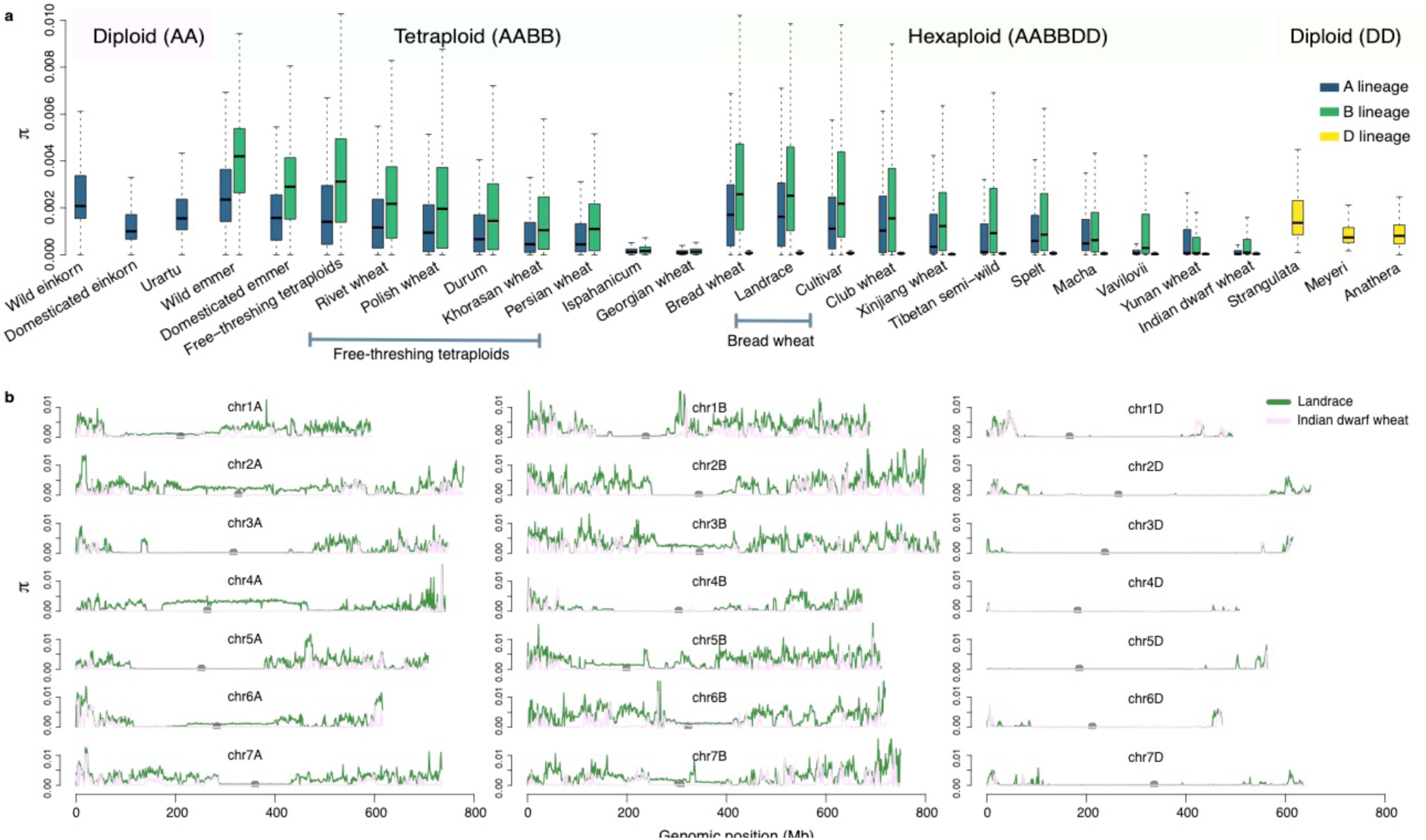
Genetic diversity of subspecies in genera *Triticum* and *Aegilops*. **a**, Diversity of all taxa by A, B and D subgenomes. **b**, Nucleotide diversity comparison between landrace and Indian dwarf wheat. Green lines represent bread wheat and pink lines represent Indian dwarf wheat.

This survey confirmed earlier studies showing that bread wheat has asymmetric distribution of nucleotide diversity on ABD subgenomes with *π*_B_ > *π*_A_ ≫ *π*D^9,11,22,23^. We provided a whole-genome estimate that *π*_D_ was only 12% and 8% of *π*_A_ and *π*_B_ (Fig. 3a and Supplementary Fig. 6a). Our results also identified the tetraploid ancestry of *π*_B_ > *π*_A_, given all subspecies of AB lineage showed the same asymmetry on AB subgenomes. The extremely low level of *π*_D_ in bread wheat could be explained by the severe bottleneck of hexaploidization^9^, since 7-fold more nucleotide diversity was observed in strangulata. In contrast, the same bottleneck seemed to have had a minor effect on genetic diversity of AB subgenomes of bread wheat, where the capture of 77% of nucleotide diversity of free-threshing tetraploids occurred. What could be the reason?

### Alien introgression into bread wheat

Recent studies identified that the interspecies introgression from wild emmer was the primary reason for the relatively high diversity of AB subgenomes in bread wheat^10,11^. However, given many complex taxonomic groups within genera *Triticum*^12,13^ (Fig. 2), the wild emmer introgression is probably only a tip of iceberg of all gene flow. Also, wild emmer introgression may bring undesired traits into bread wheat, e.g., brittle rachis and hulled grain. Therefore, there may be other more physiologically similar taxa influencing on the natural genetic diversity of bread wheat.

To test this hypothesis, we used ABBA-BABA statistics ^27,28^, which evaluate the excess of shared derived alleles between taxa, to detect gene flow from potential sources to bread wheat (Fig. 4a). Indian dwarf wheat mentioned previously presented the lowest nucleotide diversity among all hexaploids (Fig. 3a), exhibiting only 17%, 26%, and 48% of nucleotide diversity in its A, B, and D subgenomes when compared with bread wheat (Fig. 3b). An earlier report concluded that D subgenome of hexaploids preserved the phylogenetic structure of hexaploids at the dawn of their formation, because natural hybridization between hexaploids and *Ae. tauschii* is difficult^20^. It was observed that Indian dwarf wheat was basal to 96% of bread wheat in the phylogeny of D lineage (Fig. 2e). Therefore, the extremely low diversity of Indian dwarf wheat is likely to be the result of its early migration to remote areas at the southwest of the Himalayas and a consequential escape from alien introgressions. By identifying the geographically isolated and low-diversity taxon as the reference population (P1), and combining the robust phylogenies we reconstructed, comprehensive tests were enabled to detect gene flow from all species/subspecies within *Triticum* and *Aegilops*, which was not possible for previous studies^10,11^ (Fig. 4b).

**Fig. 4.**
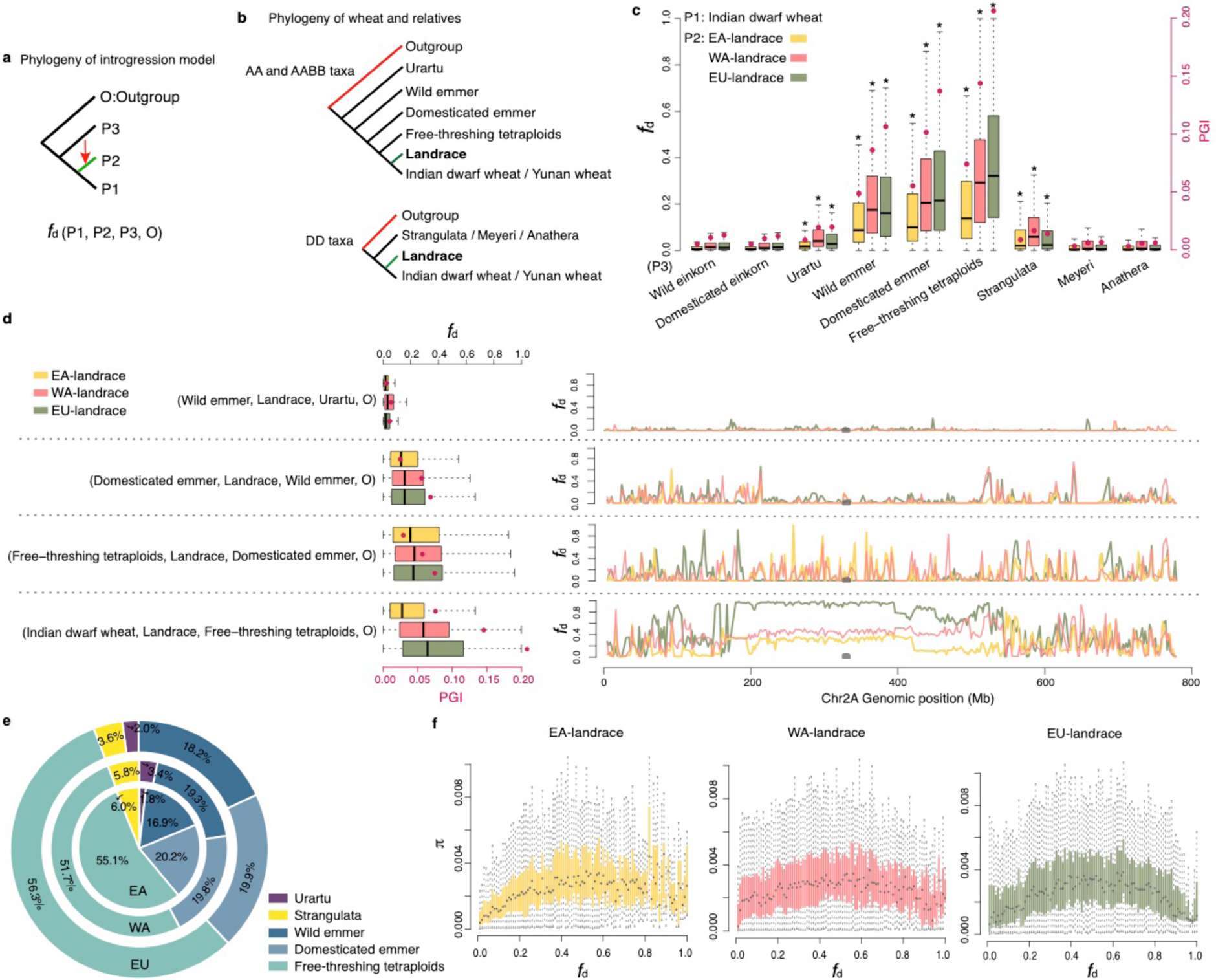
Composite introgression of *Triticum* species into bread wheat. **a**, The four-taxon topology used for modeling introgression in ABBA-BABA tests. **b**, The phylogenetic topology used for inferring introgression to landraces. Landraces were used as recipient population (P2). **c**, Gene flow from a specific donor to bread wheat using Indian dwarf wheat as P1. Star means gene flow (as indicated by *f*_*d*_)) from a certain donor was significantly higher than from wild einkorn (one-tailed *t*-test, *P* < 0.01). **d**, Taxon-specific gene flow from various donors to landraces using a cascade of P1. The picture on the left is Introgression from 4 pools to landrace across the chromosome 2A genome. **e**, The relative proportion of introgression from different donors to landraces. **f**, Relationship between nucleotide diversity (*π*) and introgression (*f*_*d*_).

Two major ABBA-BABA models to identify genomic regions of introgression were evaluated, however, following earlier assessments^28^, the *D* statistic^27^ showed inflated value for regions of low-diversity (Supplementary Fig. 7). Hence, the more stable statistic, *f*_*d*_ ^28^, which signifies gene flow when 0 < *f*_*d*_ < 1, was used to calculate the fraction of introgression in bread wheat in 100-SNP windows. Considering gene flow into bread wheat may vary by geographic regions, landraces from East Asia (EA), West Asia (WA), and Europe (EU) were tested separately. Cultivars were excluded from the analysis, since they were genetically similar to EU landraces (Fig. 2b,c).

As wild einkorn was genetically distant to bread wheat (Fig. 2a) and wild einkorn gene flow has not been reported so far, we took wild einkorn as a null and arbitrarily assumed that there was not wild einkorn gene flow happened during the evolution of bread wheat. Compared with wild einkorn, three taxa, including urartu, tetraploids, and strangulata, showed significantly higher *f*_*d*_ values (one-tailed *t*-test, *P* < 0.001, Supplementary Table 5) across the genome. Since the number of detected introgression region varied across tests, we calculated the proportion of genome of introgression (PGI) to reflect the actual genome size of bread wheat deriving from alien introgression. Both *f*_*d*_ and PGI statistics indicated that tetraploids contributed a substantial portion of the genome of bread wheat, and free-threshing tetraploids showed the highest amount of introgression to bread wheat among all taxa (Fig. 4c and Supplementary Fig. 8a). Similar to Indian dwarf wheat, Yunan wheat (*T. aestivum*. ssp. *yunna-nense*) was also a geographically isolated and low-diversity taxon (Supplementary Fig. 6b). Our results were robust to the choice of the reference population, as replacing Indian dwarf wheat with Yunan wheat produced similar outcomes (Supplementary Fig. 8b).

Shared haplotypes in tetraploids and urartu might lead to the overestimation of alien introgression to bread wheat. Therefore, we fitted the *f*_*d*_ and PGI calculation into a phylogenetic cascade to obtain PGI contributed by a specific taxon (Fig. 4d and Supplementary Fig. 9 and 10). Collectively, about 13%∼36% (EA: 13%, WA: 27%, EU:36%) of bread wheat genome come from introgression of urartu, tetraploids, and strangulata. The relatively high proportion of introgression in WA and EU landraces could be explained by their sympatric distribution with these alien donors^2^. Compared with diploids, tetraploids dominated alien introgression to bread wheat. Among them, free-threshing tetraploids were the most important source, which accounted for 52%∼56% of genomic regions of overall introgression (Fig. 4e). The largest introgression fragment from free-threshing tetraploids was on chromosome 2A (164Mb-540Mb) of the bread wheat genome, covering ∼54.5% of the whole chromosome (Fig. 4d, Supplementary Fig. 9 and 11).

Introgression did increase the genetic diversity of wheat (Fig. 4f and Supplementary Fig. 12). Taking free-threshing tetraploids introgression into EU landraces as an example, a 2.2-fold increase of nucleotide diversity was observed in genomic regions of frequent introgression (*f*_*d*_ *>* 0.01, *π =* 0.0029) compared with those of rare introgression (*f*_*d*_ *<* 0.01, *π =* 0.0013). Furthermore, we found that introgression had a non-linear effect on nucleotide diversity, of which the maximum nucleotide diversity was achieved when introgression alleles had a frequency of 0.5 in bread wheat population.

Despite the severe genetic bottlenecks of hexaploidization and domestication, the genetic diversity of bread wheat was largely compensated by composite gene flow from multiple groups of AABB taxa, including free-threshing tetraploids, domesticated emmer, and wild emmer (Fig. 5). We found 66% and 57% of nucleotide diversity of wild emmer were captured in landraces and cultivars of bread wheat, respectively. In contrast, bread wheat preserved only 14% of nucleotide diversity of its D subgenome donor, strangulata, which led to the extremely low level of nucleotide diversity in the D subgenome of bread wheat.

**Fig. 5.**
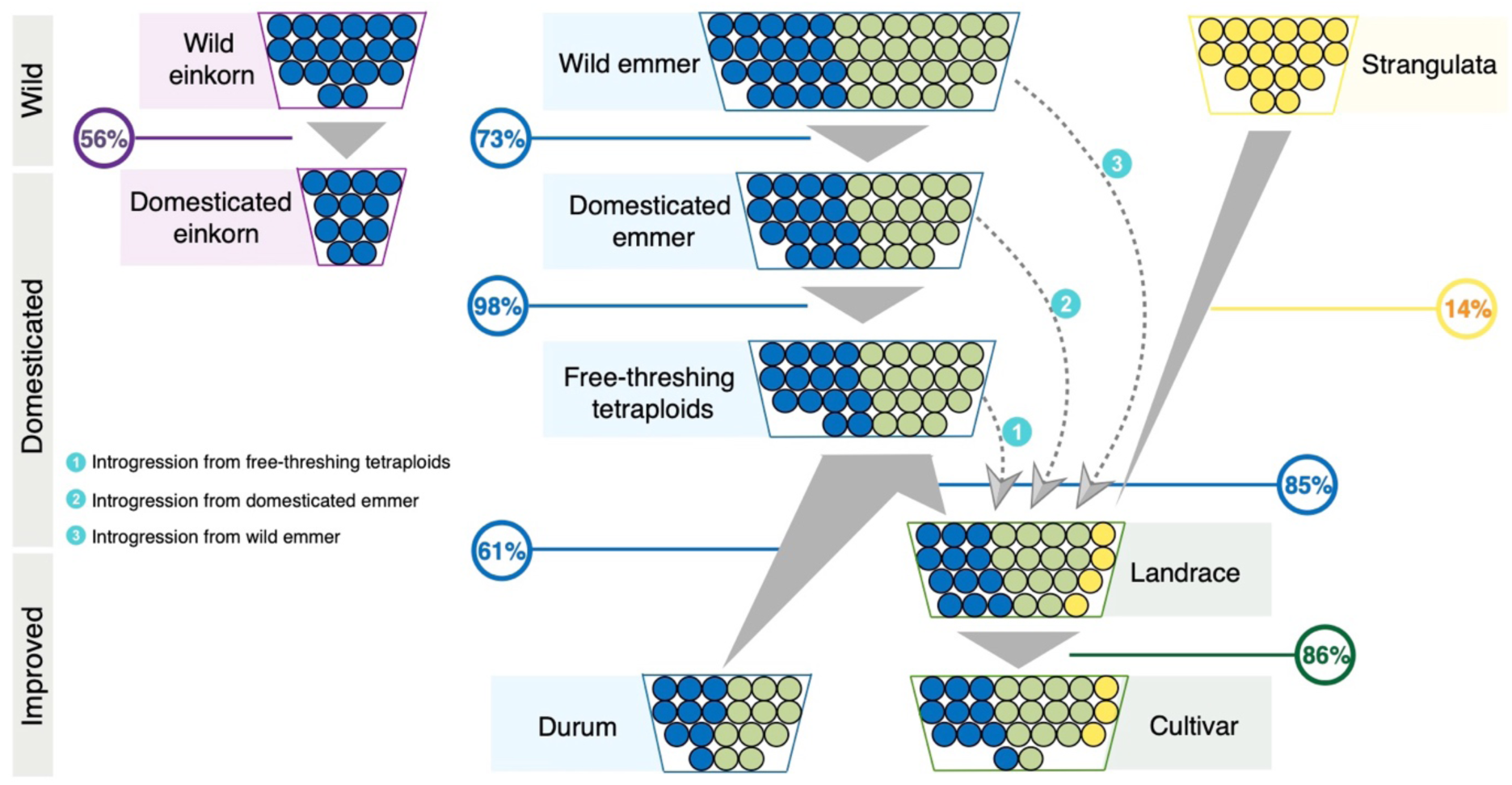
The shift of genetic diversity of *Triticum* species. The upper layer represents the wild species/subspecies, the middle layer represents domesticated species/subspecies, and the bottom layer is improved species/subspecies. Circles in a box represent nucleotide diversity of a taxon with A, B and D subgenomes represented by blue, green and yellow colors, respectively. Arrows refer to introgression from tetraploid wheats to landrace. The percentage of genetic diversity preserved in descendants is labeled in circles.

### Genomic signatures of convergent adaptation of wheats

Introgression enabled the divergent adaptation of bread wheat to global environments by substantially increasing its genetic diversity. Meanwhile, evolutionary convergence by human selection may also provide a channel of constraints that shape adaptation of bread wheat. As mentioned, at least two pairs of breeding selection events happened during *Triticum* evolution, in which wild einkorn and wild emmer were independently domesticated. Likewise, durum and bread wheat were separately improved (Fig. 1 and 6a,b). Given the vast difference in ploidy level and geographic distribution, the selected genes shared between these paired events may be of vital importance for the transition of wheats from wild grasses to major crops.

**Fig. 6.**
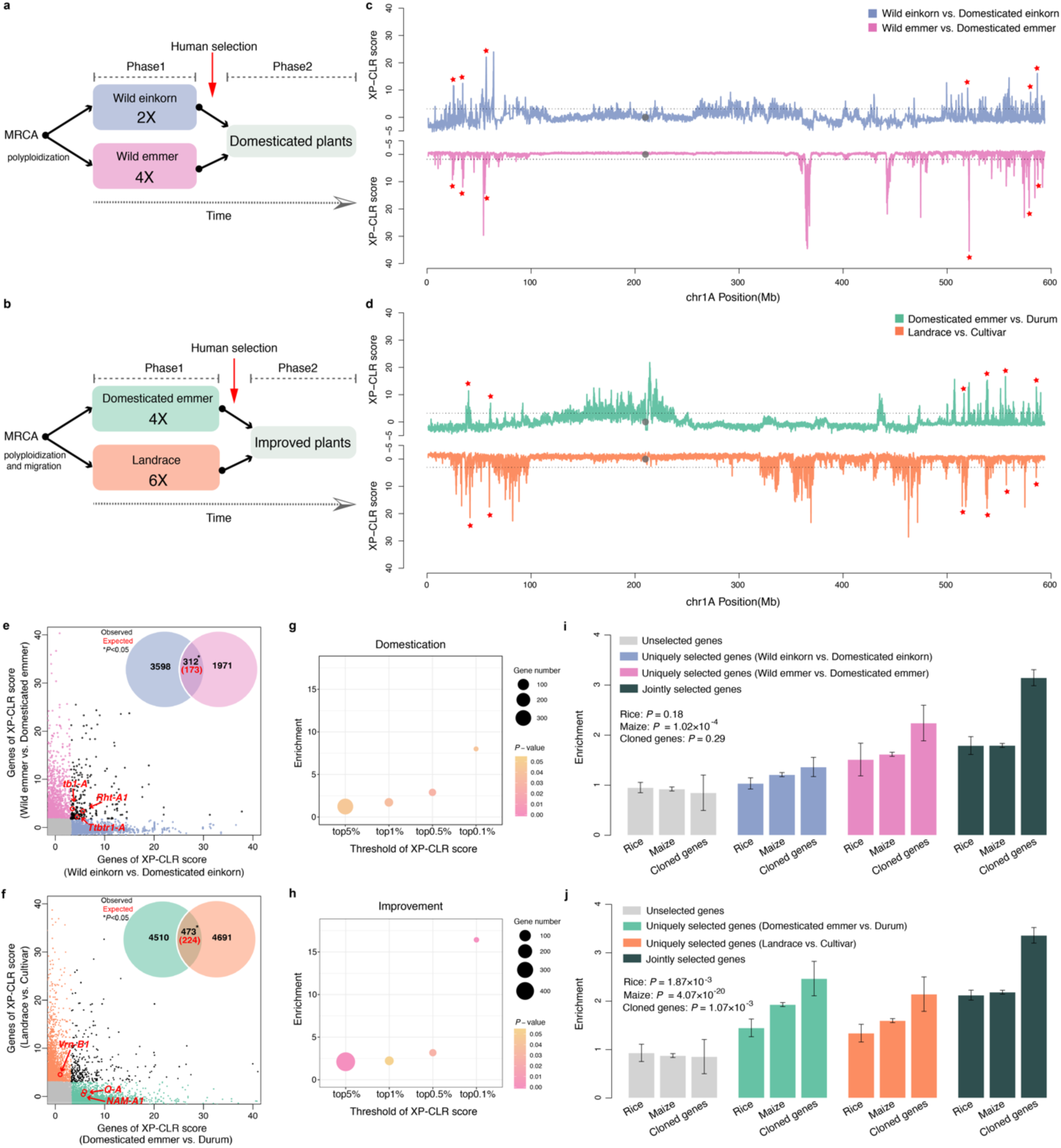
Convergent adaptation of wheats to human selection. **a and b**, The Model of domestication and improvement process. Human selection acting on the different types of plants may result in similar phenotypes. MRCA: most recent common ancestor. **c**, Selective sweeps on chromosome 1A of the paired domestication events. Top: wild einkorn vs. domesticated einkorn. Bottom: wild emmer vs. domesticated emmer. **d**, Selective sweeps on chromosome 1A of the paired improvement events. Top: domesticated emmer vs. durum. Bottom: landrace vs. cultivar. Top 20 signals with coexisting sweeps within 2 Mb intervals in corresponding selection events are labeled with red stars. **e and f**, Jointly selected genes in the paired selection events. Several cloned agronomically relevant genes are labeled. Venn diagrams show the number of jointly selected genes. Numbers in black are observed numbers under selection and numbers in red are the expected numbers under permutation test conditions. **g and h**, Enrichment for adaptive convergence under different thresholds of XP-CLR scores of the paired selection events. Circle size represents the number of jointly selected genes and circle color represents significant levels. **i and j**, Genes in selective sweeps of other crops and cloned agronomically relevant genes are enriched for selected genes in the paired selection events of wheats. *P* values represent the significance of Chi-square test.

We used the cross-population composite likelihood ratio^29^ (XP-CLR) to detect whole-genome selection signatures in the paired domestication events (wild einkorn vs. domesticated einkorn, and wild emmer vs. domesticated emmer). With the top 5% of XP-CLR scores, 3,105 and 16,141 selective sweeps were found in the genome of domesticated einkorn and domesticated emmer, respectively. About 15% of the sweeps in one selection event showed coexisting sweeps in the other event within 2 Mb intervals, suggesting convergent evolution of domestication of einkorn and emmer may have occurred (Fig. 6c,d).

However, the coexisting sweep regions might also arise from coincidence. If domestication indeed resulted in convergent adaptation of einkorn and emmer, the number of jointly selected genes of two domestication events should be much more than expected under random conditions. Within 35,345 genes in A subgenome, 3,910 (11%) and 2,283 (6.5%) genes were in the selective sweeps of domesticated einkorn and domesticated emmer. Based on the assumption of random selection of genes, 173 genes would be jointly selected (See method). However, we identified a 1.8-fold enrichment of jointly selected genes of two domestication events (*n* = 312, Permutation test, *P* = 0.047, Fig. 6e). Moreover, we found genes under strong selection during domestication were more likely to be selected in the other domestication event. By raising the threshold of XP-CLR scores from the top 5%, 1%, 0.5%, to the top 0.1%, the enrichment value increased from 1.8-fold, 2.4-fold, 3.2-fold, to 8.0-fold, despite the rapid drop in the number of jointly selected genes (Fig. 6g and Supplementary Fig. 13).

Additionally, a negative control experiment was conducted to verify the results further. As the differences between wild einkorn and wild emmer populations were caused by natural selection rather than human selection, the differentially selected genes between the two groups should not show enriched overlap with genes under selection of domestication. Performing genome-wide XP-CLR scan between wild einkorn and wild emmer, we did not observe any significant enrichment and the enrichment value dropped when increasing the threshold of XP-CLR scores (Supplementary Fig. 14 and Supplementary Table 6). Both lines of evidence suggested that the convergent evolution of the domestication of einkorn and emmer had occurred.

In addition to the analysis of paired domestication events, we also investigated the paired improvement events (domesticated emmer vs. durum, and landraces vs. cultivars) to examine convergent adaptation of wheats to recent breeding selection. Analysis done to the domestication process was also performed for the paired improvement events. Although specific values varied, e.g., number of jointly selected genes and enrichment, the same pattern of adaptive convergence was clearly exhibited (Fig. 6d,f,h and Supplementary Fig. 15). These results indicated that convergent adaptation of wheats to recent breeding had occurred as well.

Many species in the Poaceae family are cultivated as cereals to provide carbohydrates for humans. Therefore, convergent adaptation to breeding selection may be universal in cereal crops. To test the hypothesis, we used publicly available data, and then investigated whether wheat orthologs of agronomically relevant genes from other cereals were more likely to be selected in wheat breeding selection. Three data sets were used in the tests, including 1,250 genes within selective sweeps of rice domestication and improvement^30^, 3,040 genes within selective sweeps of maize domestication and improvement^31^, and 92 cloned agronomically relevant genes from rice^32^, maize^33^, and barley^34^. We identified 1,668, 5,085, and 215 wheat orthologous genes from the three data sets. They were enriched for the selected genes, especially for the jointly selected genes during wheat domestication events, rather than evenly distributed between selected genes and unselected genes (Fig. 6i). The analysis of paired improvement events (domesticated emmer vs. durum, and landraces vs. cultivars) exhibited very similar results with slightly higher enrichment (Fig. 6j). The results suggested that breeding selection has resulted in adaptive convergence in cereal crops.

### The biological function of genes showing adaptive convergence

The jointly selected genes during domestication and improvement were expected to be very important to agronomic traits of wheats. Gene ontology (GO) analysis showed that the 312 jointly selected genes in the paired domestication events were enriched for cell wall functions and glucan metabolic processes (Supplementary Fig. 16a), which could be explained by the rise of erect growth habit during early vegetative growth phase in domesticated wheats and thus requiring the strengthened stalk^35^. Meanwhile, the 473 jointly selected genes showed enrichment mostly in stimulus-response and metabolic process (Supplementary Fig. 16b), possibly reflecting recent breeding intensification efforts for disease resistance and yield.

### Convergent evolution of brittle rachis trait

The evolution of brittle rachis (BR) trait makes the most prominent example to illustrate the adaptive convergence of wheats in response to human selection. Non-BR is one of a few crucial wheat domestication traits. Independent mutations in orthologs of gene *Btr1*^36^, which controls shattering phenotype in barley, resulted in the shift from BR to non-BR in wheats, despite nearly 1 million years of divergence^6^ and different ploidy levels. In domesticated emmer, the non-BR phenotype was conferred by loss-of-function alleles in genes *TtBtr1-A* and *TtBtr1-B* in A subgenome and B subgenome, respectively^37^. Whereas in domesticated einkorn, non-BR phenotype was caused by another loss-of-function allele, a nonsynonymous mutation, within *TmBtr1-A*^38^ (Fig. 7a).

**Fig. 7.**
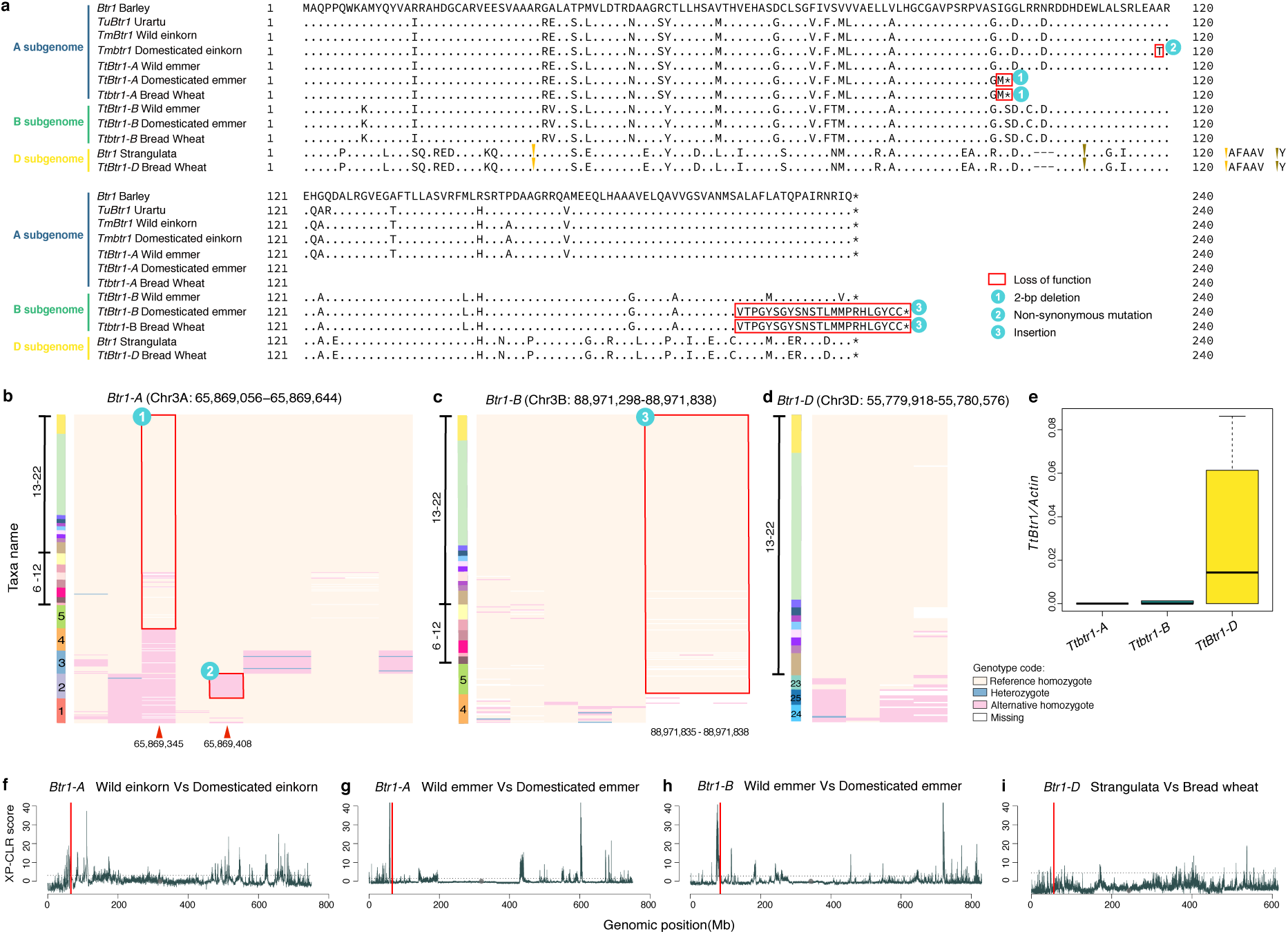
Convergent evolution of gene *Btr1*. **a**, Comparison of the amino acid sequence of orthologs of *Btr1*. Red rectangles in sequence represent loss-of-function alleles. Yellow and brown arrows are insertions, with the inserted sequence labelled at the end. An asterisk represents a stop codon. The three loss-of-function alleles are labeled with numbers from 1 to 3. **b**, Haplotypes of *TtBtr*-A in A lineage. **c**, Haplotypes of *TtBtr*-B in AB lineage. **d**, Haplotypes of *TtBtr*-D in D lineage. Haplotypes were constructed using SNPs from VMap I. Colored bars, and numbers on the left of (b, c, and d) represent different species/subspecies as shown in Fig. 2. Red arrows represent the loss-of-function alleles and the values below represent its physical position in the reference genome. **e**, Relative expression of orthologs of *Btr1* with Actin gene in spikelet of bread wheat. **f**, Selective sweeps of *Btr1*-A in domesticated einkorn. **g**, Selective sweeps of *Btr1*-A in domesticated emmer. **h**, Selective sweeps of *Btr1*-B in domesticated emmer. **i**, Selective sweeps of *Btr1*-D in bread wheat. Red lines in f to i indicate orthologs of *Btr1* gene.

We found that the three loss-of-function alleles dominated domesticated einkorn, domesticated emmer, and its descendant, hexaploid wheats (Fig. 7b,c). However, as a species with typical non-BR phenotype, bread wheat did not show any loss-of-function alleles in *TtBtr1-D*, the ortholog of *Btr1* in the D subgenome (Fig. 7a,d). As *Btr1* is expressed in spike^36^, we conducted gene expression profiling of *TtBtr1-A, TtBtr1-B*, and *TtBtr1-D* in spike of bread wheat. It turned out that the dysfunction of *TtBtr1-D* was not required for non-BR phenotype in bread wheat (Fig. 7e).

Selective sweeps identified from the domestication of emmer and einkorn exhibited strong signals of selection around *TtBtr1-A, TtBtr1-B*, and *TmBtr1-A* (Fig. 7f,g,h). Among them, *TtBtr1-A* and *TmBtr1-A* were in the list of jointly selected genes between the paired domestication events (Fig. 6e). In contrast, we did not find a selective sweep around the gene *TrBtr1-D* (Fig. 7i). Hence, the parallel fixation of loss-of-function alleles of *Btr1* orthologs in domesticated einkorn, domesticated emmer, and bread wheat demonstrated strong convergent adaptation of wheats to breeding selection.

## Discussion

Climate change and growing population are putting global food supply at risk. A business-as-usual approach alone to the genetic dissection of agronomic traits, such as quantitative trait locus (QTL) mapping and genome wide association studies (GWAS), is probably not fast enough to help fill the food gap by 2050^39^, given the limited window of time. Evolution is the ultimate field trial with proven success – it is the driving force that has integrated the various types of beneficial traits to allow plant adaptation and colonization, such as disease resistance, heat tolerance, and water use efficiency among others. Although evolution is a long-term experiment, fortunately, these successes are well preserved in the DNA sequences of extant species. Fast and effective access to the information may pave a new way for more efficient genetic study and breeding of crops^40^.

In this study, we took wheat as a model and performed the first genus-level whole-genome sequencing analysis in crops. The comprehensive sampling of *Triticum* species and the ultrahigh marker density ensured a robust phylogeny of *Triticum*. A reliable reconstruction of wheat evolutionary history will be extremely useful for many kinds of genetic research of wheat. By comparing the genetic diversity of taxonomic groups, we found that the genetic bottleneck of bread wheat resulted from polyploidization and domestication was largely compensated by a composite gene flow from multiple groups of tetraploid wheat, especially the free-threshing tetraploids. About 13%∼36% of bread wheat genome was directly contributed by introgressions. The amount of introgression in wheat genome is much higher than it in other species. For example, the human genome has only 1%∼3% introgression from Neanderthals^41^, the rice and highland maize genomes have <6% and 2%∼10% introgressions from their wild ancestors, respectively^42,43^. The high level of gene flow to bread wheat was probably due to the substantially different effective population size (*N*_*e*_) between two taxa as indicated by the scenario of introgression in maize^44^. Also, it could be related to the polyploid speciation of hexaploid wheats, where high level of reproductive isolation occurred immediately between bread wheat and its genetic donors. The increase of genetic diversity from alien introgression is fundamental to the global expansion of bread wheat, indicating the importance of introducing genetic diversity into current wheat cultivars.

Apropos of wheats, it is hard to ignore the brilliance of their recent adaptation, the success in both diverse global environments and the uniform “table environment.” By comparing paired selection events (both domestication and recent improvement) of *Triticum* species, we identified significant footprints of convergent adaptation to breeding selection in wheats, where the enrichment of adaptive convergence ranges from 1.8-fold to 16.4-fold relative to expected value at random conditions, despite the drastic difference in ploidy levels and growing zones. The jointly selected genes were enriched for cell wall functions, stimulus-response, and metabolic processes, meaning that these genes, screened by nature’s experiments, can be useful to breed cultivars for lodging-resistance, stress-smartness, and high-yields. The non-brittle rachis trait demonstrated convergent evolution of wheats well. Also, it should be noted that only a few genes showed high-level convergence in terms of fold enrichment. Given the strong selection for carbohydrates by humans, the incomplete convergence may come from differences in environmental conditions and target traits, genes with redundant functions, complex genetic architecture of yield and its inherent linkage drag, or merely random drift by large. Perhaps, 10,000 years of human selection is not long enough for these annual crops to reach their fitness peak in the agricultural context. However, the very incomplete convergence also suggests a great potential to combine adaptation success across species for crop improvement. Leveraging the advancement in gene editing technologies^45^, speedy test of genes showing adaptation success within *Triticum* species and even other plants is anticipated to expedite genetic studies and breeding of bread wheat.

## Acknowledgements

We thank Dr. Yalong Guo from Institute of Botany, Chinese Academy of Sciences, Dr. Zhiyong Liu, and Dr. Shusong Zheng from Institute of Genetics and Developmental Biology, Chinese Academy of Sciences, and Dr. Philip Kear from International Potato Center (CIP) - China Center for Asia and the Pacific for their valuable suggestions and comments. This research was supported by Strategic Priority Research Program of the Chinese Academy of Sciences (XDA24020201) and the National Natural Science Foundation of China (31970631) to F.L., the National Natural Science Foundation of China (31921005) to Y.J., and China Postdoctoral Science Foundation (2018M631614) to Y.Z.

## Author contributions

Y.Z. and X.Z. performed the experiments with assistance from J.X., H.C., Y.W., Y.-G.W., A.B., J.W. and S.L.; Y.L. and C.Y. helped collect the materials; Y.Z., X.Z., F.L., C.J., H.L., and S.L. analyzed the data. F.L and Y.J. designed and supervised the research. F.L., Y.Z. and X.Z. wrote the manuscripts. All authors discussed the results and commented on the manuscript.

## Competing interests

The authors declare no competing interests.

## Data availability

The raw sequence data have been deposited in Sequence Read Archive (https://www.ncbi.nlm.nih.gov/sra) under accession PRJNA439156 and Genome Sequence Archive (https://bigd.big.ac.cn/gsa) under accession numbers PRJCA001501. The genotype data in VMap I will be public available as soon as the manuscript is published.

## Methods

### Plant material and planting

To represent high diversity in wheat, we collected a total of 414 accessions from 71 countries worldwide. This population contains 25 species/subspecies, covering all subspecies of AA, AABB and AABBDD genomes in *Triticum* and two subspecies of DD genome in *Aegilops tauschii* (Supplementary Table 1 and 2). Plant materials are available at the National BioResource Project (NBRP), Genebank Gatersleben of Leibniz Institute of Plant Genetics and Crop Plant Research (IPK), National Small Grains Collection (NSGC), International Centre for Agricultural Research in Dry Areas-Syria (ICARDA), and Chinese Crop Germplasm Resource Information System (CGRIS).

Seeds of each accession were first sterilized in 30% H_2_O_2_ for 10 minutes, then thoroughly washed for five times with distilled water. The sterilized seeds were germinated in water at 25 °C for 2 days. After germination, seeds with endosperm residue were transplanted into soil and grown in greenhouse. After a 15-day to 30-day growth under long-day conditions and a 2-day etiolation processing, the leaves were sampled.

### Sequencing and quality control

The whole population was sequenced in two batches. The first batch of 119 bread wheat accessions was sequenced with an average depth of 10x. The second batch of 300 accessions was sequenced with an average depth of 3.8x. The overall average sequencing coverage was ∼6x. Each accession was sequenced in either on HiSeq 2500 or NovaSeq 6000 Illumina machine using 150bp pair-end reads. All raw data were quality controlled by filtering out low quality reads and adapter pollutions. We removed the reads with more than half of bases having a quality less than Q20 or more than 5% bases with N.

### Variant calling

Given the different ploidy levels across the population, we developed a specific pipeline for variant calling (Supplementary Fig. 1). Firstly, the reference genome (IWGSC RefSeq v1.0) was divided into four different taxonomic groups based on their genome constitution, namely AA, AABB, AABBDD and DD taxa, respectively. Number of accessions contained in AA, AABB, AABBDD and DD taxa was 91, 125, 173 and 30, respectively. Then, all accessions were mapped to their corresponding reference with BWA-MEM^46^. Finally, Sentieon (http://www.sentieon.com) was used to obtain the raw variants for each group.

### Quality control of SNPs

Although all accessions in our study are inbred with low heterozygosity, the nature of polyploidy and low-coverage of depth may still cause false positive in SNP calling due to misalignment and sequencing error, we applied a specific pipeline with six filters to eliminate low confidence SNPs. The low confidence SNPs were identified by either technological or genetic filters. Technically, SNPs with low mapping quality, biased depth distribution, or potential sequencing error were considered unreliable. Genetically, SNPs broken the genetic law, such as linkage disequilibrium or identity by descent, should not be retained. The numbers of SNPs left after each filter were represented from Supplementary Table 7 to 10 for AA, AABB, AABBDD and DD taxa, respectively. The SNP filters we used are listed below.

#### Quality filter

SNPs with low mapping quality is unreliable. We firstly cleaned the SNPs based on the alignment quality. Any SNPs met one of the following situations were eliminated with custom scripts: Quality By Depth (QD) was less than 2; Fisher strand score (FS) was larger than 60; Strand Odds Ratio (SOR) value was less 3; RMS Mapping Quality (MQ) was less 30. For AA taxa, about 45% of raw SNPs were filtered out in this step, while for AABB, AABBDD and DD taxa, 27%, 25% and 30% of raw SNPs were filtered out, respectively. Totally, about 297M SNPs were filtered out for all four taxonomic groups.

#### Depth filter

To avoid the bias caused by misalignment, we removed outliers with ultra-high or low total depth. For all four taxonomic groups, SNPs on genomic regions greater or less than 30% of mode value from site depth distribution were filtered out. SNPs with high depth variance in each group were also eliminated. SNPs with standard deviation of the depth greater than 3 were filtered. For AA taxa, the filtered SNPs accounted for about 45% the remaining SNPs of the previous step. For AABB taxa, the filtered SNPs accounted for about 13% the remaining SNPs of the previous step. For AABBDD taxa, the filtered SNPs accounted for about 10% the remaining SNPs the previous step. For DD taxa, the filtered SNPs accounted for about 21% the remaining SNPs of the previous step. Finally, ∼126 million SNPs were filtered out in this step.

#### Segregation test (ST) filter

Misalignment could cause high heterozygosity, which could be eliminated based on testing whether allelic depth distribution is random or not^47^. The SNPs which *P*-value of segregation test larger than 0.01 would be taken as random and eliminated as such randomness indicating potential errors. Scripts from previous study in maize were used for filtering^48^. There were not many SNPs filtered in this step as wheat is inbred with few heterozygosity sites and the low sequencing coverage. About 1.8M SNPs were filtered out in DD taxa, the number of which was the highest among the four groups, while the number of AA, AABB and AABBDD taxa was 1.1M, 0.4M and 0.9M, respectively. Totally, about 4.2M SNPs were filtered out in this step.

#### Linkage disequilibrium (LD) filter

The true SNPs should be in the local linkage with adjacent SNPs. We calculated the squared correlation (r^2^) with the most adjacent 50 SNPs, namely 25 SNPs upstream and 25 SNPs downstream for each site. Sites with mean r^2^ less than 0.1 were eliminated. This step filtered more in polyploidy groups, as 9.4% and 5.7% SNPs in AABB and AABBDD taxa were filtered while only 3.4% and 1.7% SNPs were filtered in AA and DD taxa. Totally, about 36.6M SNPs were eliminated.

#### Identity by descent (IBD) filter

SNPs that located in the IBD region are expected to be identical and variants violated IBD constraint should be eliminated. To determine IBD regions, SNPs left in the previous step with the highest MQ value were used as anchor SNPs to derive the IBD information for each group. A total of 13,245,787; 32,034,359; 28,826,874 and 14,952,087 SNPs were used as anchors in AA, AABB, AABBDD and DD taxa, respectively. The genetic divergence between any two accessions was calculated using 2000 anchor sites^48^. Considering the mean distance across all pairs is more than 0.3 (Supplementary Table 11), two accessions were considered to be in IBD if the genetic divergence was ≤ 0.03, which was set as the same as previous described^48^. In AA taxa, numbers of taxa involved in IBD relationships ranged from 12 to 91 with an average of 83. The IBD distance ranged from 0.24 to 0.48, with an average of 0.39. The number of SNPs filtered out in this step were about 9.1M. In AABB taxa, numbers of taxa involved in IBD relationships ranged from 11 to 125 with an average of 113. The IBD distance ranged from 0.16 to 0.48, with an average of 0.32. The number of SNPs filtered out in this step were about 13.6M. In AABBDD taxa, numbers of taxa involved in IBD relationships ranged from 6 to 54 with an average of 46. And the IBD distance ranged from 0.11 to 0.52, with an average of 0.34. The number of SNPs filtered out in this step were about 3.4M. In DD taxa, numbers of taxa involved in IBD relationships ranged from 3 to 30 with an average of 27. And the IBD distance ranged from 0.20 to 0.45, with an average of 0.36. The number of SNPs filtered out in this step were about 1.6M. A total of about 36.6M SNPs were eliminated in this step.

#### Minor allele count (mac) filter

Rare SNPs could be detected with high false positive rate. Since population genetic tests in the study mostly rely on common SNPs to keep the statistical power, we eliminated rare SNPs using --mac 3 in VCFtools (0.1.15)^49^, indicating that all alleles must be detected in at least 2 different accessions. For all four taxonomic groups, the rare SNPs ratios were 30%, 38%, 31% and 26% for AA, AABB, AABBDD and DD taxa, respectively. We eliminated ∼155M SNPs in this step.

### Detection of syntenic sites

Given the fact that the taxa containing A, B, and D subgenomes have different levels of coalescence and the reference bias of SNP calling can be severe when coalescence is deep. For fair comparison between A, B, and D subgenomes in this research, we analyzed SNPs only on conserved regions / syntenic sites. We did not do genome alignment to detect syntenic sites, because the einkorn reference genome was not available. We identified the syntenic sites based on mapping depth distribution, and SNPs on syntenic sites were kept. We detected the syntenic sites by lineages as following: Firstly, mapping depth per site of reference genome was calculated using samtools v0.1.19^50^; then, we calculated the total depth and depth standard deviation across the lineage for each site using custom scripts; finally, sites passed the depth filter as described previously were defined as syntenic. We identified 657,315,587, 2,320,149,443 and 1,877,720,646 sites as syntenic on A, B and D subgenome, comprising about 13%, 45% and 48% of the whole subgenomes, respectively (Supplementary Table 12). The syntenic sites were used to filtered for syntenic SNPs of VMap I after merging SNPs as described below.

### Construction of VMap I by merging SNPs from four taxonomic groups

For the construction of VMap I, SNPs in all four taxonomic groups were merged as an union by lineages with custom scripts. For example, for A lineage, we merged all SNPs on A subgenome, including AA, AABB and AABBDD taxa. After merging and filtering for syntenic SNPs, we constructed the SNP library of the VMap I containing a total of 103,899,092 SNPs. Then, we genotyped all accessions to build the VMap I by scanning their bam files using HapMapScanner (https://github.com/PlantGeneticsLab/TIGER/tree/master/src/pgl/app/hapScanner). The genotype likelihood was calculated on the basis of a multinomial test, as previously described^51^. After genotyping, the overall missing rate was less than 6.4% in VMap I. Number of SNPs on each chromosome was represented in Supplementary Table 13. The distribution of polymorphic SNPs across different groups in A, B and D lineage were summarized from Supplementary Table 14 to 16.

### Evaluation of false discovery rate for SNP calling

To evaluate the accuracy of VMap I, we used the SNPs called by the reads of the reference genome Chinese spring as false positive control. Short sequence reads (Supplementary Table 17) used for construction of reference were sampled from each chromosome. As the reference genome was sequenced by chromosome, so we could also divide all reads into four taxonomic groups: AA taxa only include reads from A subgenome, AABB taxa includes reads from A and B subgenomes, AABBDD taxa including all reads, DD taxa only include reads from D subgenome. Variants for each group were called using the pipeline above without any filtering. We detected 514,363, 1,309,263, 928,327 and 588,006 SNPs for AA, AABB, AABBDD and DD taxa, respectively. A total of 396,468 SNPs was found to be false positives in VMap I, with the overall false positive rate of approximately 0.024%.

### Removing of samples

To avoid the potential mistakes on sampling, we estimated the alignment rate, missing rate and heterozygous rate for each sample after filtering. The alignment rate was calculated with samtools v0.1.19^50^; Missing rate and heterozygous rate were calculated with plink (v1.90b6.7)^52^. A total of 5 samples were excluded due to relative low alignment rate (1 tetraploid accession), large missing rate (2 tetraploid accession and 1 hexaploid accession) or high heterozygous rate (1 tetraploid accession). These samples were probably contaminated during sampling, or sequencing process. They are not included in the 414 samples in VMap I.

### Genetic divergence between all accessions and Chinese Spring

To evaluate the genetic distance between Chinese Spring (CS) and each accession, we estimated pair-wise genetic divergence (*d*_*xy*_) for all accessions using TASSEL5^53^. Only corresponding SNPs on subgenome in CS were used. For example, when compared with AA taxa, we only used SNPs on A subgenome for CS.

### Inference of ancestral allele

To inference the ancestral allele for each SNP, we used barley (*Hordeum vulgare*) and *Aegilops tauschii* as outgroups for A and B subgenomes, and *Hordeum vulgare* and *Triticum urartu* as outgroups for D subgenome^54,55^. NUCmer program implemented in the latest release of MUMmer4 (v4.0.0.beta2)^56^ was used to align the outgroups genome to Chinese Spring with --maxmatch -g 1000 -c 90 -l 40. Sites overlapped with VMap I were retained using custom scripts. SNPs which were not biallelic after outgroups were added were further eliminated. Only alleles identical in outgroups were taken as ancestral allele.

### Phylogenetic analysis

We randomly selected ∼200K SNPs for each phylogeny reconstruction. Phylogenetic trees of each group were reconstructed using RAxML software (v8.2.12)^57^ with a hundred times bootstrap in GTRGAMMA model, and the output tree was plotted in iTOL^58^. For A lineage, we firstly reconstructed the tree using 100 times bootstrap for all 385 accessions including barley. Barely was set as outgroup through -o parameter. The other parameters were -f a -m GTRGAMMA -p 12346 -x 12346 -# 100. For AB lineage, which contains 293 accession of all tetraploid and hexaploid wheat, we randomly sampled SNPs from both A and B subgenomes. The phylogenetic tree was reconstructed with 100 times bootstrap and wild emmer was set as outgroup through -o parameter. The other parameters were set as same as the previous analysis. For D lineage, we reconstructed the tree with 100 times bootstrap for all 203 accessions with barely as outgroup. Then we re-analyzed the D lineage by removing the two varietas from *Ae. tauschii* ssp. *tauschii*. The tree of all 181 accessions in the D lineage was reconstructed with 100 times bootstrap with the 7 strangulata accessions as outgroup. The other parameters were set as same as the previous analysis.

### Ancestry coefficient analysis

Individual ancestry coefficient for A lineage, AB lineage or D lineage was performed by sNMF (Version 1.2)^59^ with K varied from 2 to 7. The randomly sampled SNPs dataset for phylogenetic analysis were also used in this analysis. As we know the classification of each sample, the best K was determined with the smallest value which could separate the population as expected. All analysis was repeated 100 times and the results were clustered with CLUMPP^60^. Other parameters used in sNMF and CLUMPP were set by default.

### Nucleotide diversity analysis

To compare the diversity among subgenomes in species/subspecies, we calculated the nucleotide diversity for each subgenomes across the genome in a window of 10 kb with following function^61^. However, to prevent potential underestimation, only syntenic sites rather than the whole 10-kb window were used for calculation.

### Population differentiation (*F*_*ST*_)

Weir & Cockerham’s *F*_*ST*_ value per site between two populations were calculated using VCFtools (0.1.15)^49^, and values smaller than 0 was set to 0 manually when calculating the mean value. To estimate the mean *F*_*ST*_ value between two populations, we averaged the sum *F*_*ST*_ of each site by the number of syntenic sites.

### Introgression analysis with ABBA-BABA test

A four-taxon model ((P1, P2), P3, O) was used to estimate Patterson’s *D* statistic (ABBA-BABA test). The *D* statistic of the whole genome was calculated as described before^62^. If there is no introgression happened from the population P3 to P2, the expected *D* statistic would be zero. If there existed introgression from P3 to P2, the *D* statistic would be significantly larger than zero, while the significant negative value indicating introgression happed from P3 to P1. To perform the *D* statistic test, we divided the chromosome into 1Mb bins and then performed block jackknife for calculating Z-score. *D* statistic was taken as significant if the absolute Z-score was greater than 3.

### Testing whole-genome introgression to bread wheat using the *D* statistic

To estimate the introgression to bread wheat, we separated the landrace into three groups based on their geographic distribution (EA-, WA-, and EU-landrace) based on *D* statistic, we used barley as the outgroup, Indian dwarf wheat or Yunan wheat as P1 and each landrace group was used as P2. The left taxon were tested as the potential donor population (P3). The estimation was followed from https://github.com/simonhmartin/tutorials/tree/master/ABBA_BABA_whole_genome.

### Estimation of *f*_*d*_ statistic in sliding windows

The *D* statistic only reflects the whole genome introgression, in contrast, *f*_*d*_ statistic^63^ can estimate the introgression proportion in a given window. The *f*_*d*_ statistic ranges from -1 to 1, theoretically. Under a given four-taxon topology ((P1, P2), P3, O), positive *f*_*d*_ statistic value indicates introgression proportion from population P3 to P2, while zero means no introgression. Different with the *D* statistic, the negative *f*_*d*_ statistic value does not have biological meanings.

We estimated the *f*_*d*_ values across the genome using the python code available at (https://github.com/simonhmartin/genomics_general). The sliding window was set with window size of 100 SNPs and step size of 50 SNPs. The minimum good sites in each window was set to 3 through -m flag. For windows of *D* < 0, or of *D* > 0, but *f*_*d*_ > 1, the *f*_*d*_ statistic value becomes meaningless or noisy, therefore we converted the *f*_*d*_ statistic value to zero.

We estimated *f*_*d*_ statistic value with Indian dwarf wheat or Yunan wheat as P1, the three subgroups of landrace (EA-, WA- and EU-landraces) as P2. All other species/subspecies or all free-threshing tetraploid wheat were used as P3, separately.

### Relationship between introgression and nucleotide diversity

As *f*_*d*_ statistic in window was more robust than *D* statistic, it was used to evaluate the relationship between nucleotide diversity and introgression. To estimate the diversity in the same window of *f*_*d*_ statistics, we calculated the diversity in this window as described above. As the nucleotide diversity ranged from 0 to 1, we grouped the nucleotide diversity of each window into 100 non-overlap bins with a bin size of 0.01. The boxplot of *f*_*d*_ statistic for each bin was plot using R software.

### Proportion of genome of introgression (PGI)

To estimate the proportion of introgression from P3 to P2, we firstly set *f*_*d*_ statistic to 0 while *D* statistic or *f*_*d*_ statistic in this region was less than 0 or larger than 1. Proportion of genome of introgression (PGI) was defined as the overall proportion of introgression across the whole genome. For i^th^ window, *f*_*di*_ refers to the *f*_*d*_ value in this window, *G*_*i*_ refers to window size in base pair and *G* is the genome size in base pair. PGI could be estimated with following formula:

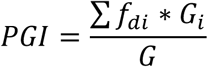

### Selection sweeps detected by XP-CLR in wheat

To detect the selection sweeps, we used XP-CLR test as it is more robust than other methods^64^. In wheat, XP-CLR score between two populations were estimated using XP-CLR software with -w1 0.005 500 10000 -p1 0.95. Genetic distance was estimated from the recombination rate data from previous publication^65^. R package GenWin was used for the normalization XP-CLR statistic and detecting the boundary of genome region with smoothness = 2000 and method = 4.

To detect which gene were under selection, we ranked the selection sweeps with XP-CLR score in descending and took the top 5%, 1%, 0.5%, and 0.1% regions as selected. Considering the larger linkage disequilibrium in wheat, we took genes as selected as long as it overlapped with selection sweeps.

### Detection of genes under convergent evolution in wheat

To detect genes showing convergent evolution, we tested the enrichment of jointly selected genes in the paired domestication events (wild einkorn vs. domesticated einkorn and wild emmer vs. domesticated emmer) and the paired improvement events (domesticated emmer vs. durum and landrace vs. cultivar). Enrichment was defined as the ratio of the observed number and expected number of jointly selected genes. The expected number of jointly selected genes was estimated under using permutation method. To evaluate the level of convergence at different selection intensity, we also raised the thresholds of XP-CLR scores from the top 5%, to 1%, 0.5%, to the top 0.1%. The significance of convergence at different levels was tested under the hypergeometric distribution test^66^ and permutation test. To general the null distribution of jointly selected genes in two parallel selection events, taking the paired domestication events for example, we shuffled accessions of wild einkorn and domesticated einkorn then resampled them into two groups to perform XP-CLR scan. We also shuffled accessions of wild emmer and domesticated emmer then resampled them into two groups to perform XP-CLR scan. Then an expected number of jointly selection genes at random conditions can be calculated by comparing their XP-CLR results. The procedure was repeated for 1000 times to yield a null distribution of the number jointly selected genes. The mean of the null was taken as the expected number of jointly selected genes between the paired domestication events.

### Selection sweeps in maize and rice

The whole genome XP-CLR results for maize were publicly available^67^, but the XP-CLR results of rice were not. Previously published data of rice (http://server.ncgr.ac.cn/RiceHap3/Genotype.php) were download from RiceHap3 database^68^, and XP-CLR test was performed with -w1 0.02 200 2000 -p1 0.95. GenWin was used with smoothness = 200 and method = 4. Genomic region with XP-CLR scores ranked the top 5% were defined as selection sweep regions. There were 150 regions under selection and 1250 genes were located in those selection sweep regions.

### Identification of wheat orthologs

To identify wheat orthologous genes with rice, maize and barley, we combined all wheat genes (IWGSC RefSeq annotation v1.1), genes with selective sweeps in rice, domestication and improvement relevant genes reported in maize^67^, and also domestication relevant genes in barley^69^. The OrthoMCL pipeline^70^ was applied to compute the all-against-all similarities with BLASTP E value < 1 × 10^−5^. If there were multiple orthologs in wheat one subgenome, we only kept the best BLASTP hit as candidate.

### Gene ontology analysis

GO annotation terms for *Triticum aestivum* genes (IWGSC RefSeq annotation v1.1) were downloaded from Ensembl Plants Genes 44 database (https://plants.ensembl.org/biomart/martview/). GO analysis was performed with agriGO (version 2.0)^71^. Enrichment significance was analyzed with the Fisher test. Enrichment results with more than 5 annotations and FDR< 0.05 were plotted with R packages clusterProfiler (version 3.10.0).

### Identification of the wheat orthologs to the barley *Btr1* and *Btr2* genes

To identify the exact position of *Btr1* and *Btr2* orthologs in wheat, *Btr1* and *Btr2* gene sequences were BLAST to the reference genome of wheat (IWGSC RefSeq v1.0), and hits were found on the short arms of chromosomes 3A, 3B and 3D (Supplementary Table 18). *Btr1* and *Btr2* genes orthologs were found on chromosome 3A (designated *TtBtr1-A* and *TtBtr2-A*, respectively), chromosome 3B (designated *TtBtr1-B* and *TtBtr2-B*, respectively), and chromosome 3D (designated *TtBtr1-D* and *TtBtr2-D*, respectively), alongside 3-4 duplicated copies for each gene. We identified the specific location of *TtBtr1* in the three subgenomes based on BLAST identity and confirmed by comparing the distances and directions of these copies with wild emmer. The *TtBtr1-A* gene position is 3A:65,869,035 - 65,869,715 bp, the *TtBtr1-B* gene position is 3B:88,971,271-88,971,836 bp and *TtBtr1-D* gene position is 3D:55,780,576-55,779,918 bp. The *TtBtr2-A* gene position is 3A:65,829,549 - 65,828,837 bp, the *TtBtr2-B* gene position is 3B:87,718,581-87,717,921bp and *TtBtr2-D* gene position is 3D:56,282,328-56,281,668 bp.

### Relative expression of *Btr1* of bread wheat

As *Btr1* is expressed in spike, we conducted gene expression analysis of *TtBtr1-A, TtBtr1-B* and *TtBtr1-D* in spike of bread wheat. Chinese Spring spike transcriptome data (SRR6799266, SRR6799267, SRR6799268, SRR6802609, SRR6802610, SRR6802611, SRR6802614) were download directly from NCBI SRA database. Gene expression was quantified using kallisto^72^ with standard settings. Due to *Btr1* was not included in IWGSC RefSeq annotation v1.1, we added the annotation to dataset manually.

Althougth *TtBtr1-A, TtBtr1-B* and *TtBtr1-D* all have a low expression level compared with the house-keeping gene *Actin* (*TraesCS5A02G274400*), *TtBtr1-D* was expressed relative higher than *TtBtr1-A* and *TtBtr1-B*. That might mean *TtBtr1-D* was not required for non-BR phenotype in bread wheat.

### Identification of loss-of-function alleles in the *TtBtr1* genes

To identify the mutations in the *TtBtr1*, we constructed its haplotype of all accessions. The domesticated einkorn features a point mutation which introduces a non-synonymous allele at 359 from alanine to threonine. In domesticated emmer, free-threshing tetraploids and all hexaploids, the *Ttbtr1-A* allele features a 2 bp deletion (290 bases from the start codon) that introduces a premature stop codon, thereby truncating the wild type 196 AA protein to only 97 AA; the *Ttbtr1-B* allele features an insertion (539 bases from start codon) that results in a longer C terminus protein sequence compared to the wild type 196 AA protein. These mutations are expected to disrupt the structures of the *Ttbtr1-A* and *Ttbtr1-B* proteins and are therefore considered to be loss-of-function alleles. No polymorphisms were detected within the coding regions of the *Ttbtr1-D* protein.

