## Supplementary Figures for "Convergence within divergence: insights of wheat adaptation from *Triticum* population sequencing"

### Title

[5]Annoroad Gene Technology Corporation, Beijing, China.

[6]Novogene Bioinformatics Institute, Beijing, China.

[7]CAS-JIC Centre of Excellence for Plant and Microbial Science (CEPAMS), Institute of Genetics and Developmental Biology, Chinese Academy of Sciences, Beijing, China.

### Author list footnotes

[#]These authors contributed equally to this work.

[*]Corresponding authors

### Correspondence

 (F.L.); (Y.J.)

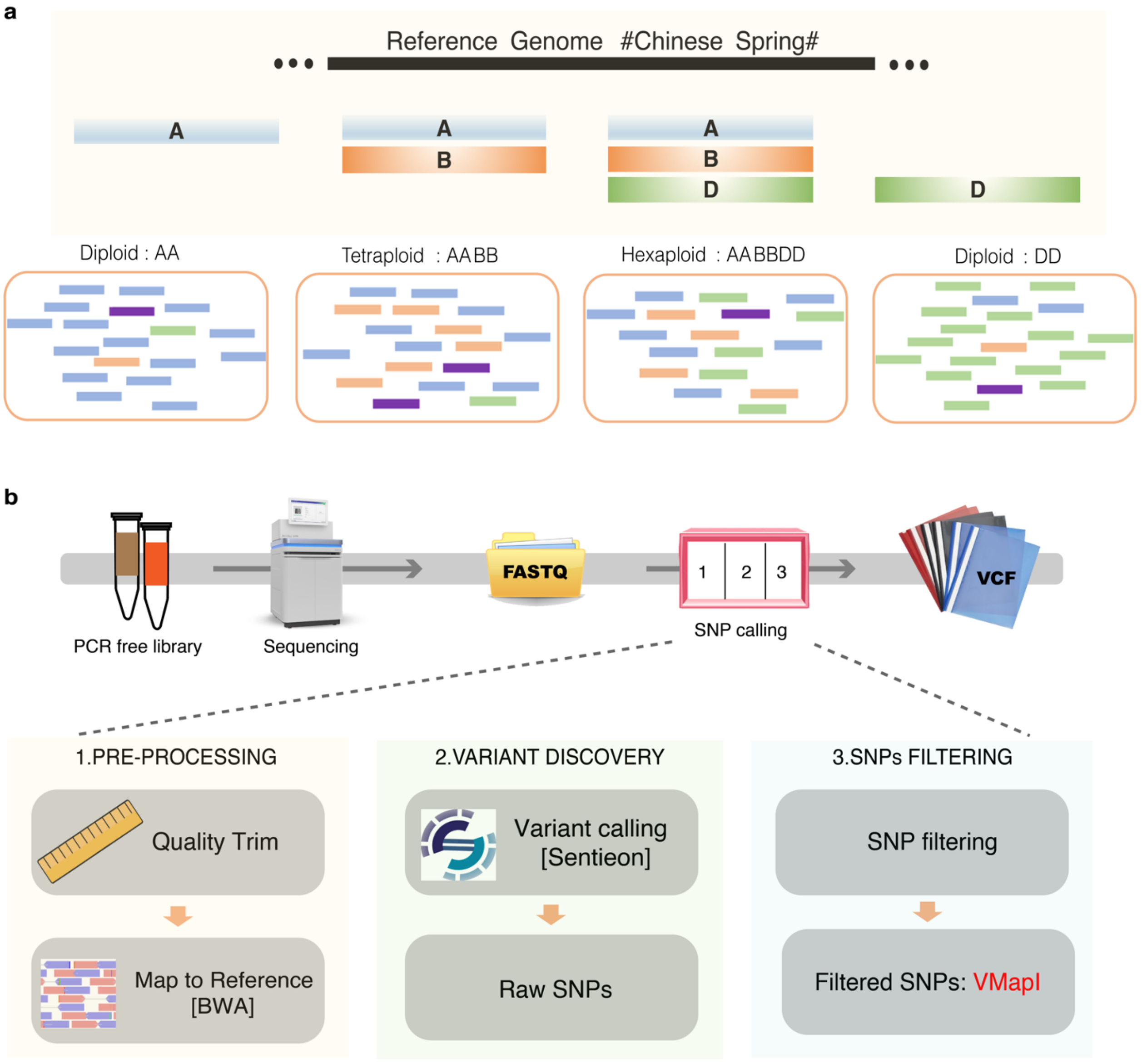


**Supplementary Fig. 1.** Pipeline of cross-ploidy variation discovery. **a**, Reference genome of genome/subgenome groups. The reference genome was divided into four taxonomic group based on population genome constitution (AA, AABB, AABBDD and DD). **b,** Variation calling pipeline. We built PCR free library of all samples, which were then sequenced on Illumina machine. SNPs were called using BWA and Sentieon softwares. After filtering raw SNPs, all SNPs were merged by lineage.

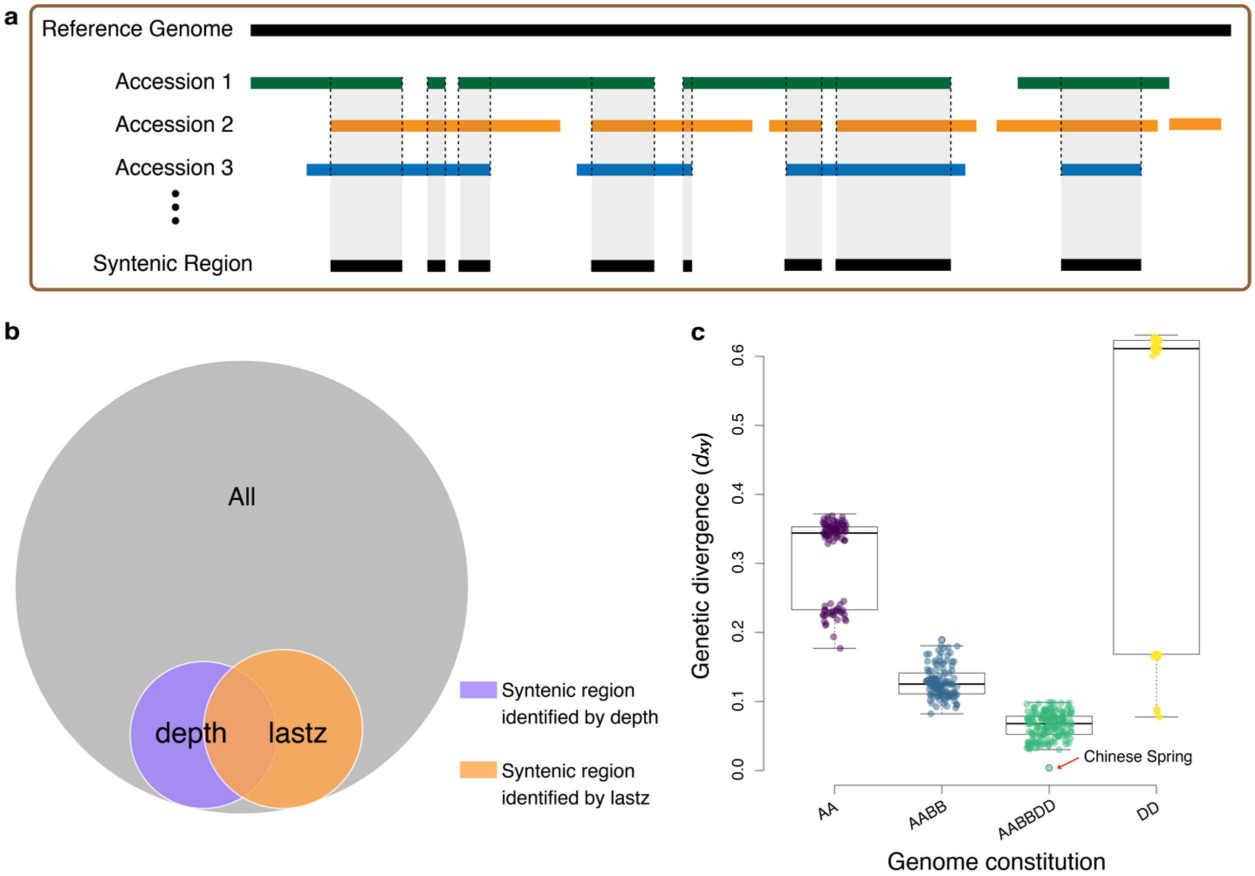


**Supplementary Fig. 2.** Evaluation of variation calling. **a**, Definition of syntenic region. A site whose depth passed the depth filter is defined as syntenic. Genetic variations in syntenic regions of each lineage were retained. **b,** Comparison of syntenic sites identified by depth and lastz methods. The gray circle represents all sites in the chromosome, the purple circle represents the syntenic sites defined by depth-based method and the yellow circle represents the syntenic sites defined by lastz. To verify syntenic sites detected by the depth approach, we also detected the syntenic sites using whole-genome alignment to examine the consistency between the two approaches. We aligned the wild emmer genome[1] to the reference with lastz and the sites with unique mapping were defined as lastz-defined syntenic sites[2]. For example, we detected the syntenic sites on the second part of chromosome 1A with overall length of 122,798,051 bp, we totally detected 13,061,714 and 15,383,247 syntenic sites using depth-based and lastz, respectively, and 5,445,193 sites were detected as syntenic using both methods. We found that 41.69% of depth-defined syntenic sites were also lastz-defined syntenic sites. The two approaches showed consistency, but the overlapping of syntenic sites is not high. Given the prevalent structural variations in plant genomes, the depth approach based on alignment of hundreds of genomes is likely to be more sensitive and accurate on detecting syntenic sites than aligning only two single genomes. **c,** The genetic divergence ($d_{xy}$) between reference genome and each accession was estimated. Chinese Spring was used as a calibration accession and had the lowest $d_{xy}$ (0.0037) as indicated by the red arrow in AABBDD taxa.

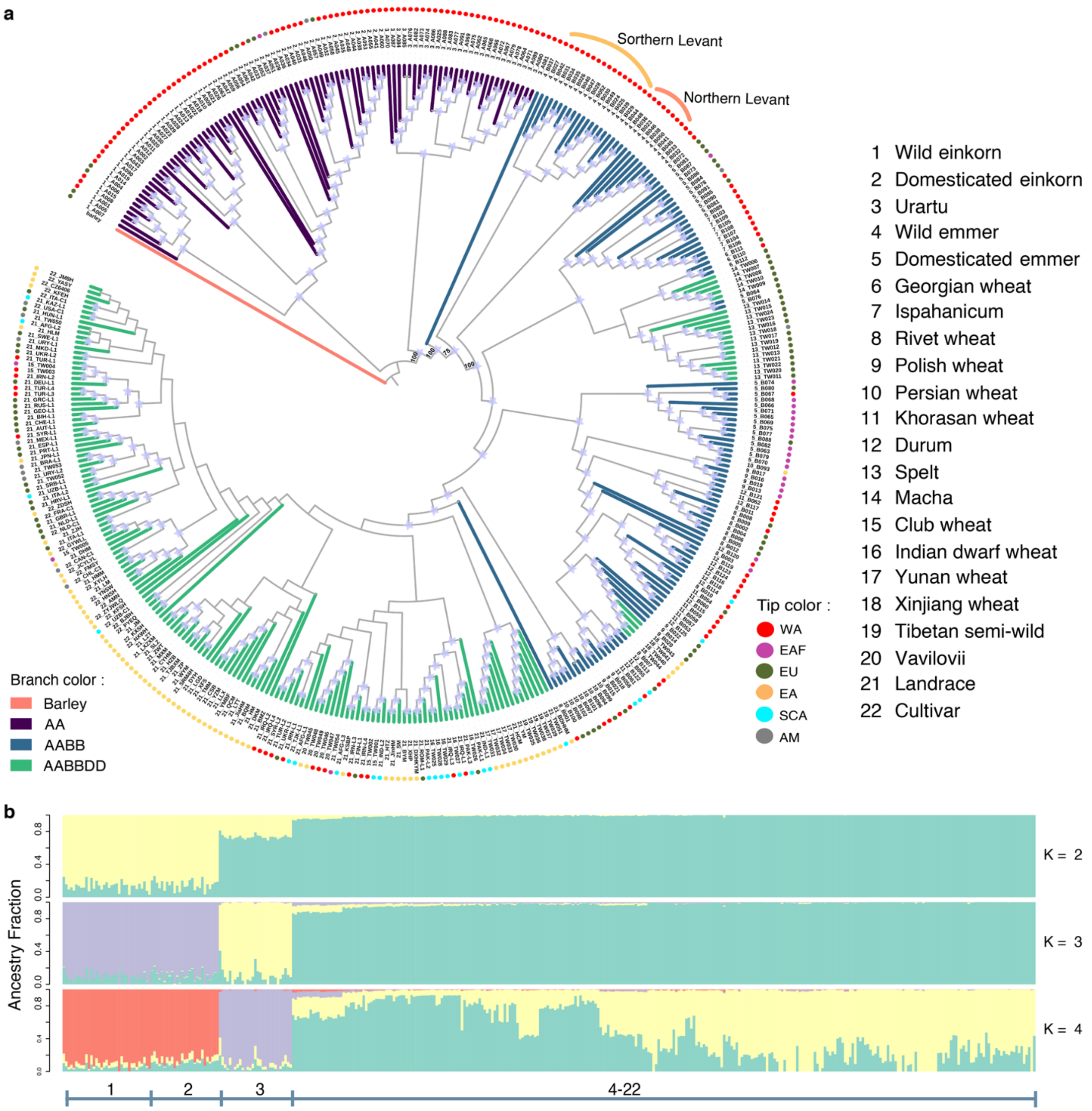


**Supplementary Fig. 3.** Population structure of A lineage. **a**, Phylogenetic tree of A lineage. Barley was used as the outgroup. Species/subspecies were represented by numbers from 1 to 22. Tip colors of the phylogeny represent sampling region of each accession, while branch colors represent ploidy levels. Branches with reliable bootstrap value (> 50) are labeled with a purple pentagram at corresponding nodes. WA: West Asia; EAF: East Africa; EU: Europe; EA: East Asia; SCA: South and Center Asia; AM: America. **b,** Ancestry coefficient analysis of A lineage with K = 4. Species are labeled with their corresponding numbers and colors.

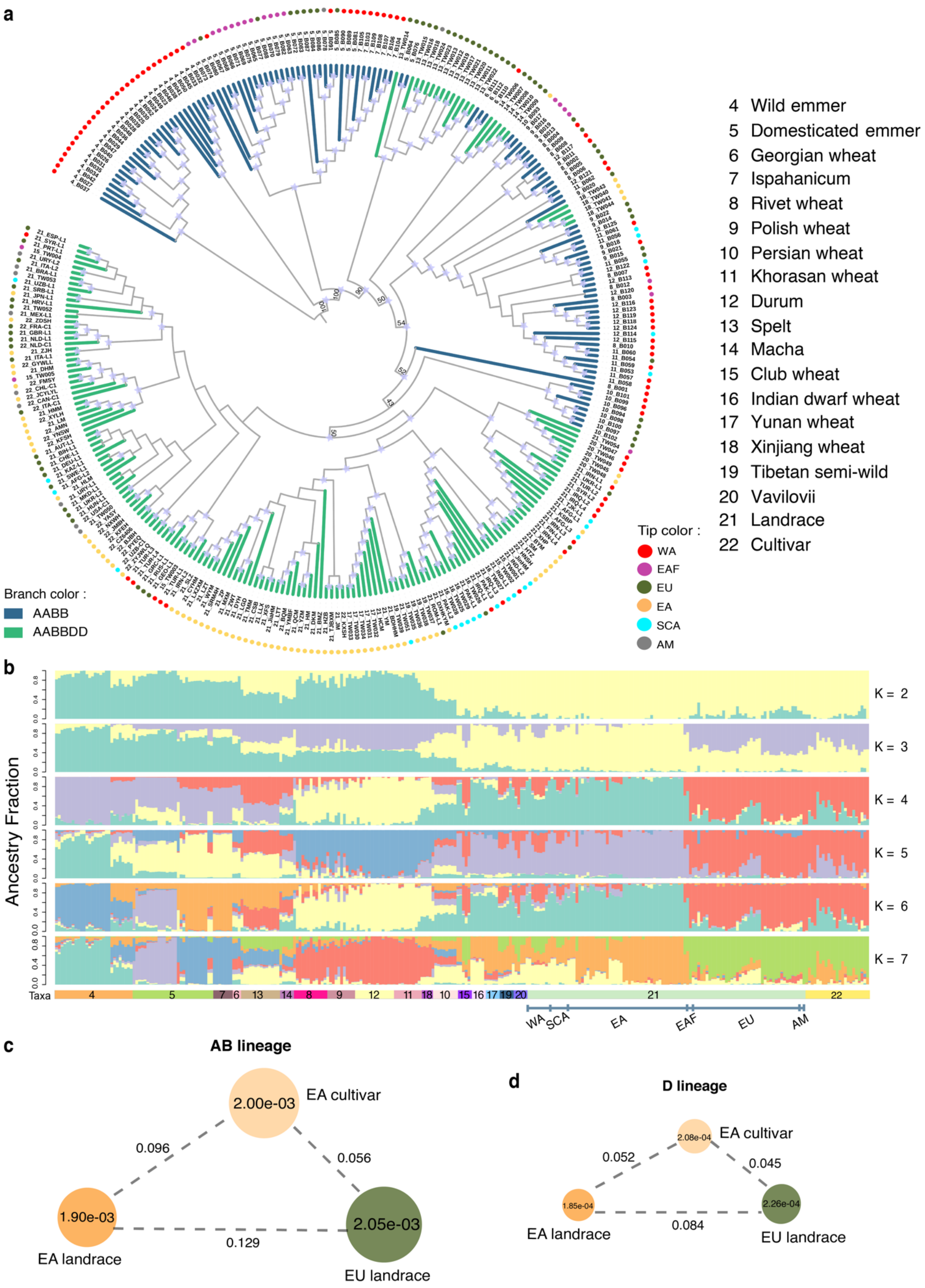
a
4 Wild emmer
5 Domesticated emmer
6 Georgian wheat
7 Ispahanicum
8 Rivet wheat
9 Polish wheat
10 Persian wheat
11 Khorasan wheat
12 Durum
13 Spelt
14 Macha
15 Club wheat
16 Indian dwarf wheat
17 Yunan wheat
18 Xinjiang wheat
19 Tibetan semi-wild
20 Vavilovii
21 Landrace
22 Cultivar
Tip color :
WA
EAF
EU
EA
SCA
AM
Branch color :
AABB
AABBDD
b
Ancestry Fraction
K = 2
K = 3
K = 4
K = 5
K = 6
K = 7
Taxa
WA
SCA
EA
EAF
EU
AM
c
AB lineage
2.00e-03
EA cultivar
0.096
0.056
1.90e-03
2.05e-03
0.129
EA landrace
EU landrace
d
D lineage
2.08e-04
EA cultivar
0.052
0.045
1.85e-04
2.26e-04
0.084
EA landrace
EU landrace

**Supplementary Fig. 4.** Population structure of AB lineage. **a,** Phylogenetic tree of AB lineage. Wild emmer from southern levant was used as the outgroup. Species/subspecies were represented by numbers from 1 to 22. Tip colors of the phylogeny represent sampling region of each accession, while branch colors represent ploidy levels. Branches with reliable bootstrap value (> 50) are labeled with a purple pentagram at corresponding nodes. WA: West Asia; EAF: East Africa; EU: Europe; EA: East Asia; SCA: South and Center Asia; AM: America. **b,** Ancestral coefficient of AB lineage. K varied from 2 to 7. Species are labeled with their corresponding numbers and colors. **c,** Nucleotide diversity and population differentiation ($F_{ST}$) in AB lineage. **d,** Nucleotide diversity and population differentiation ($F_{ST}$) in D lineage. Values in the circle represent nucleotide diversity and numbers next to dashed lines represent $F_{ST}$.

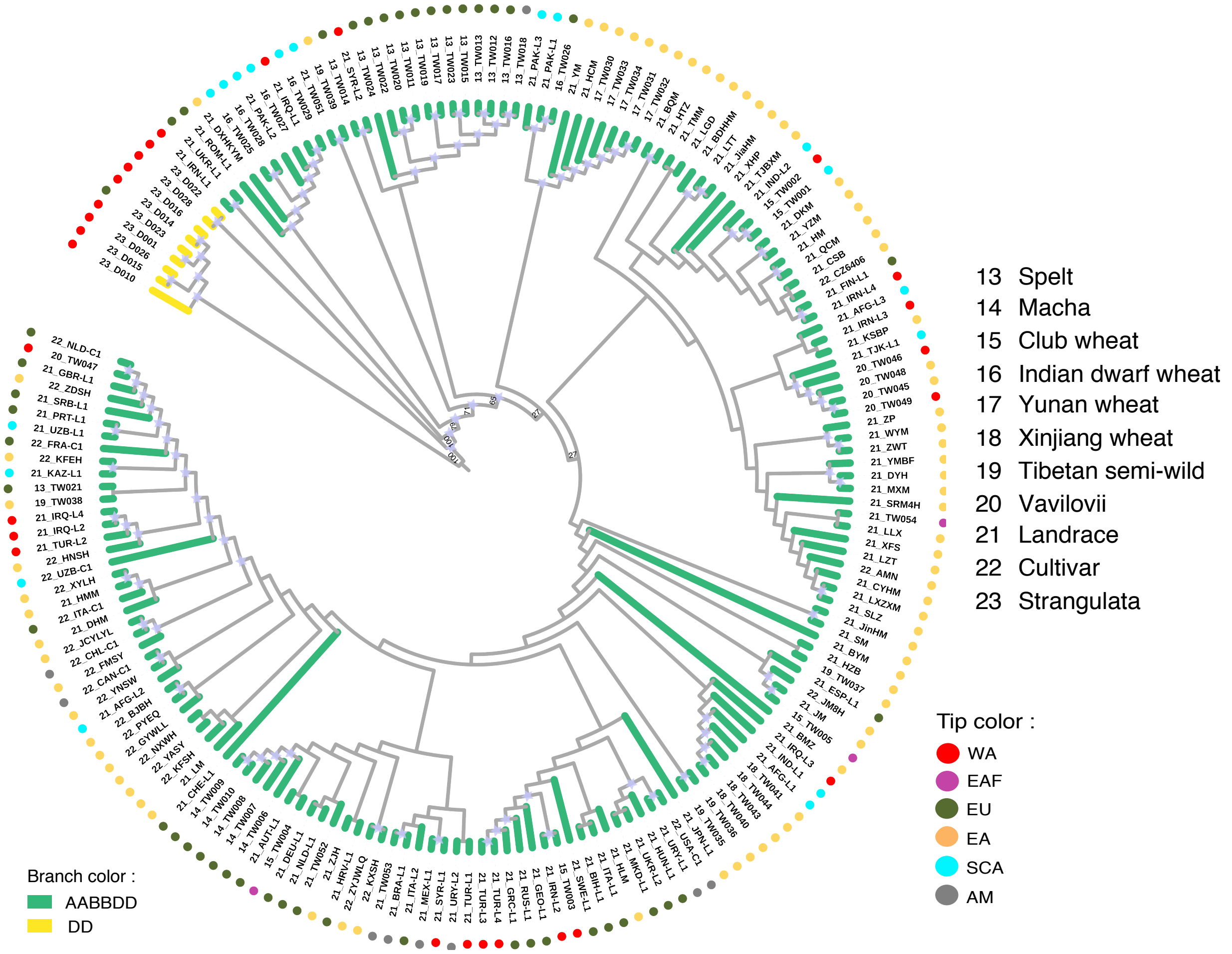


**Supplementary Fig. 5.** Phylogenetic tree of D lineage. Strangulata was used as the outgroup of the phylogeny of D lineage. Species/subspecies are represented by numbers from 1 to 25. Tip colors of the phylogeny represent sampling region of each accession, while branch colors represent ploidy levels. Branches with reliable bootstrap value (> 50) are labeled with a purple pentagram at corresponding nodes.

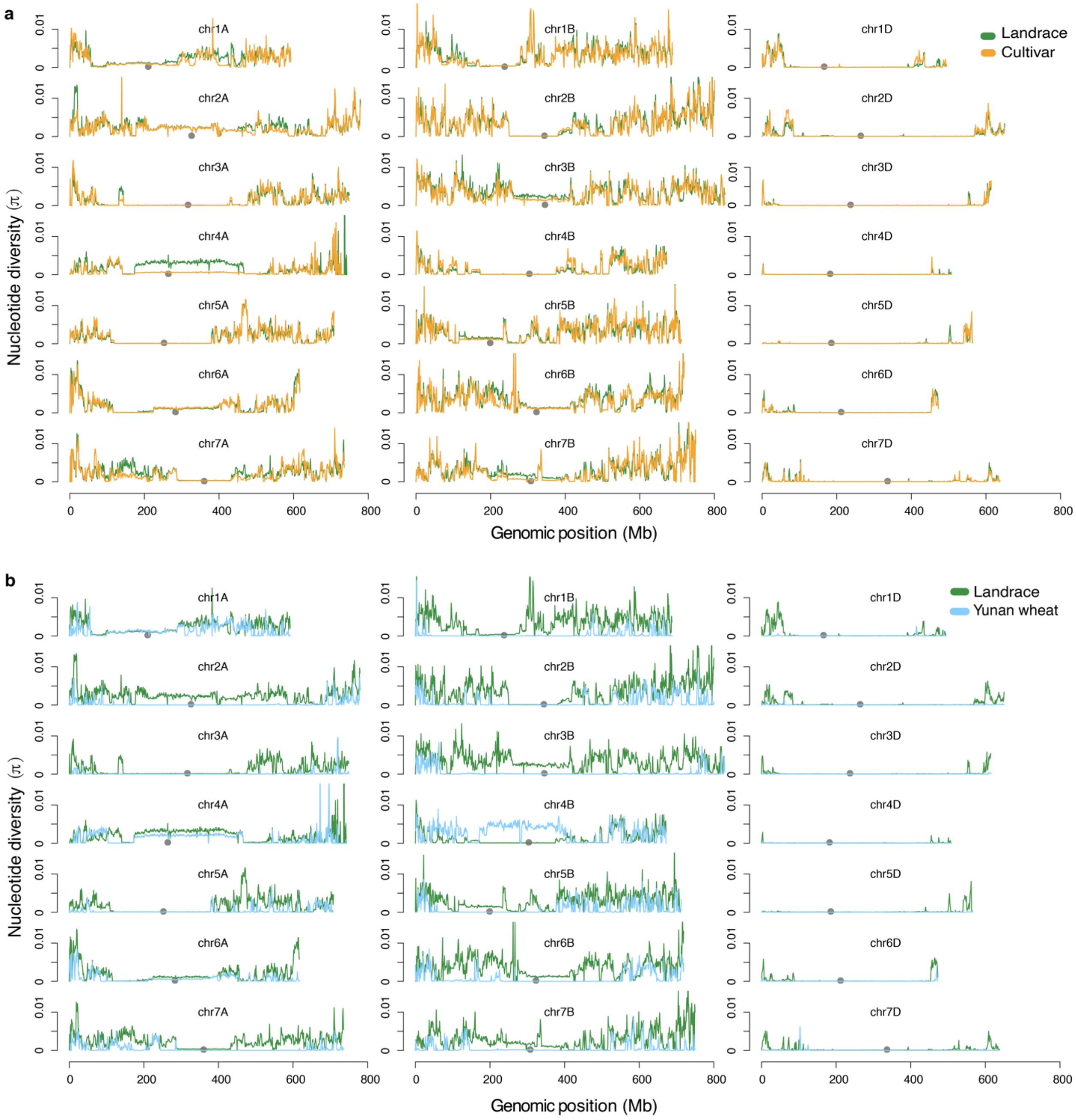


**Supplementary Fig. 6.** Comparison of nucleotide diversity of wheats across the genome. **a,** Comparison of nucleotide diversity of bread wheat across each chromosome. **b,** Comparison of nucleotide diversity of landrace and Indian dwarf wheat across the genome. Green lines represent landrace, yellow lines represent cultivar and blue lines represent Yunan wheat. Grey dots are the chromosome centromeres.

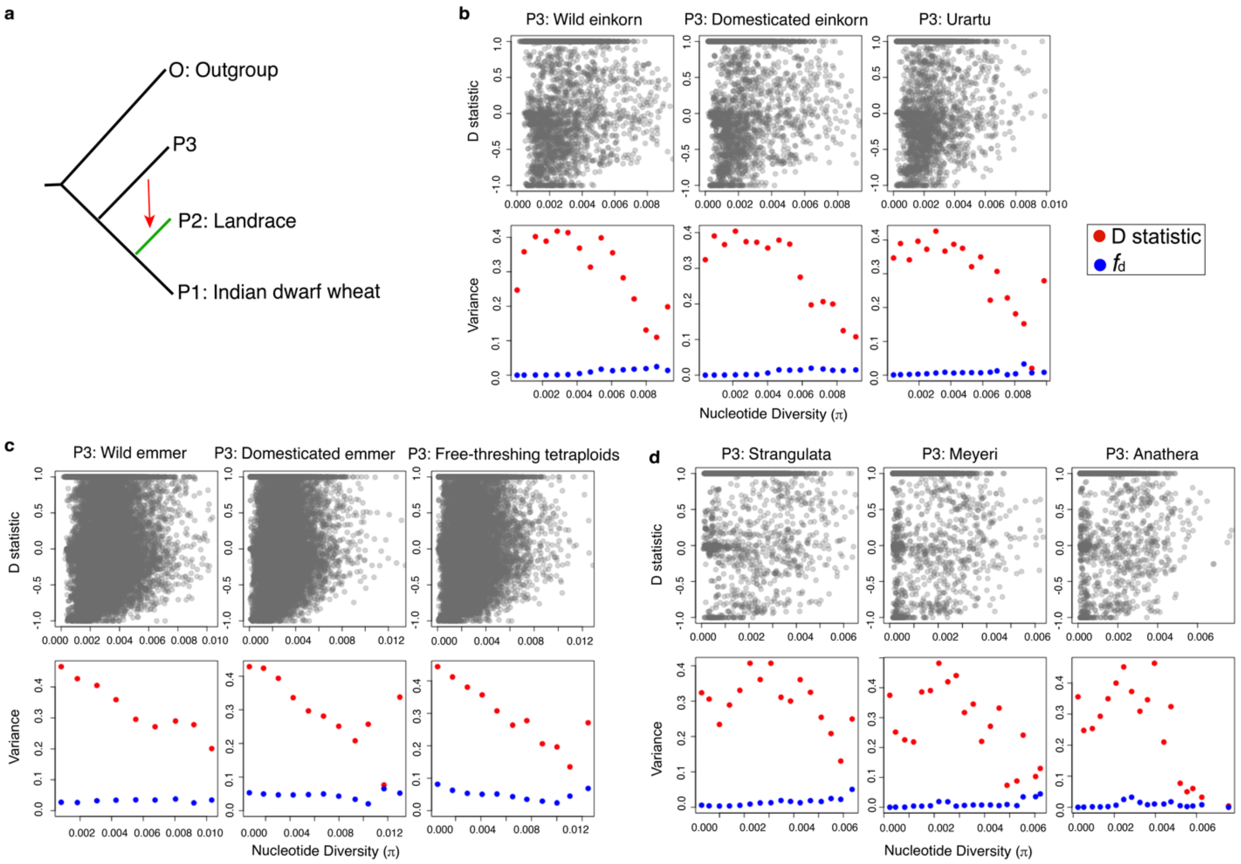


**Supplementary Fig. 7.** Bias of *D* statistic at low-diversity genomic regions. **a,** Four-taxon topology for estimating *D* statistic and $f_d$ with Indian dwarf wheat as P1 and European (EU) landraces as P2. **b,** *D* statistic (top), variance of *D* statistic and $f_d$ (bottom) as a function of nucleotide diversity in A lineage. **c,** *D* statistic (top), variance of *D* statistic and $f_d$ (bottom) as a function of nucleotide diversity in AB lineage. **d,** *D* statistic (top), variance of *D* statistic and $f_d$ (bottom) as a function of nucleotide diversity in D lineage. Red dots represent variance of *D* statistic and blue dots represent variance of $f_d$.

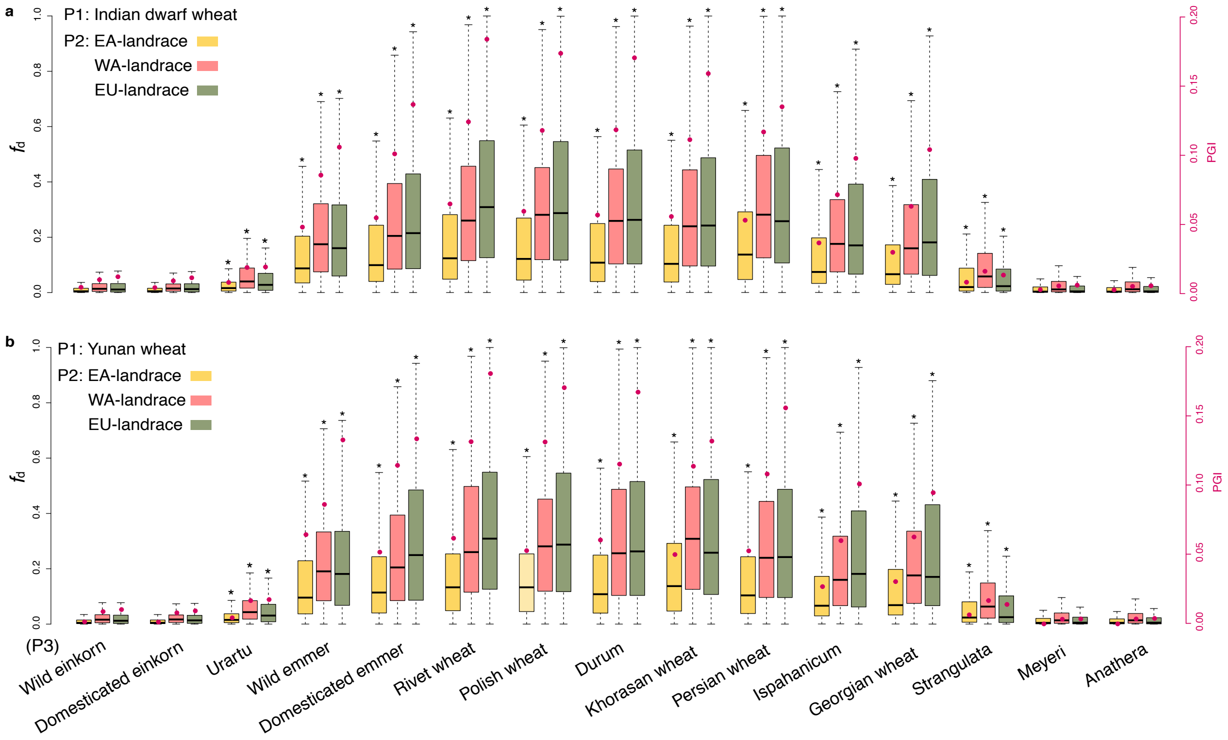


**Supplementary Fig. 8.** The Proportion of introgression estimated from different species/subspecies to landrace. **a,** Introgression estimated with Indian dwarf wheat as P1. **b,** Introgression estimated with Yunan wheat as P1. $f_d$ statistic (left y-axis) was estimated under four-taxon topology ((P1, P2), P3, O). Indian dwarf wheat and each subgroup of landraces were used as P1 and P2, respectively. Each P3 group is shown in x-axis. Right y-axis shows PGI statistic from different species/subspecies.

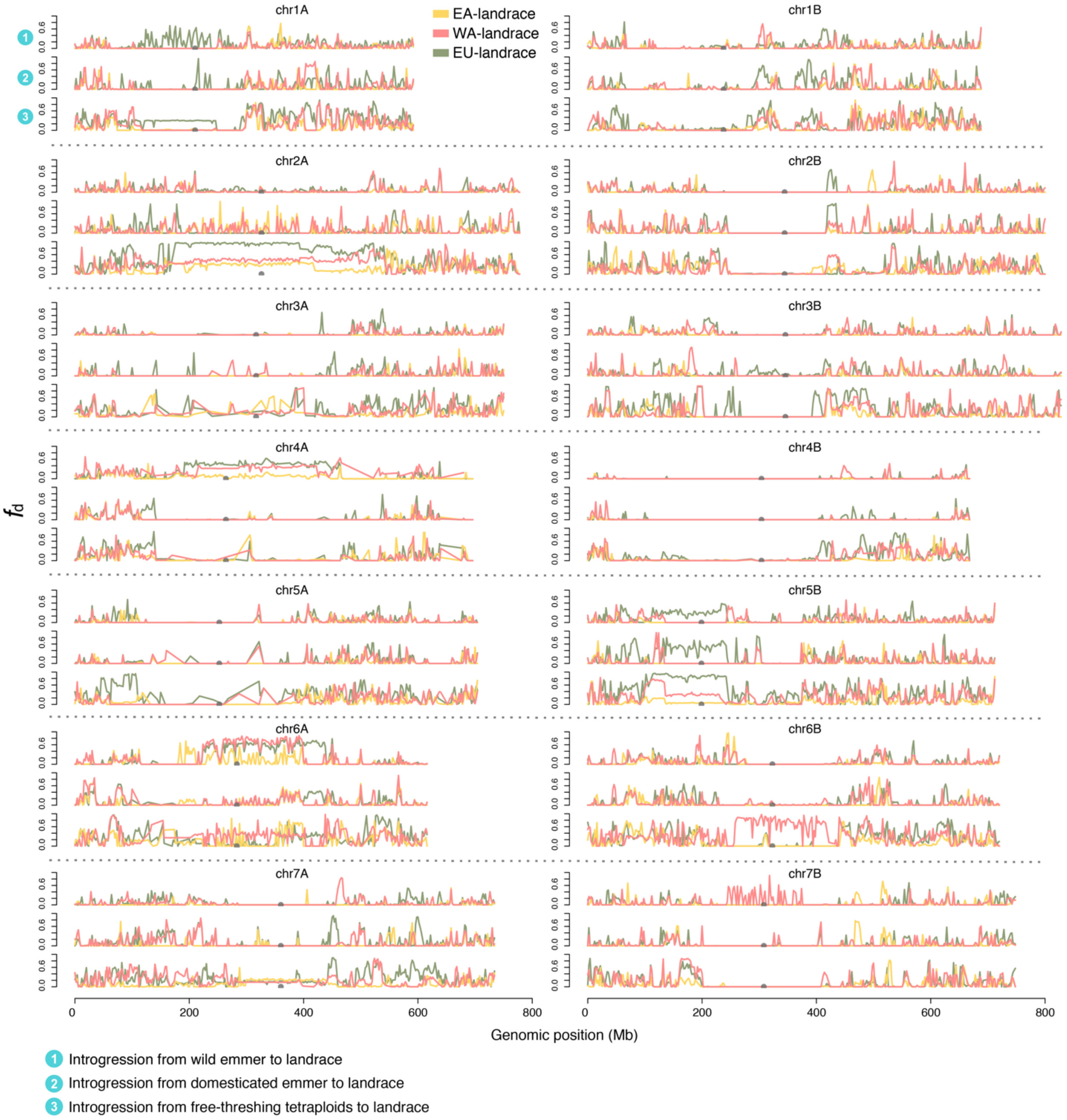


**Supplementary Fig. 9.** Introgression from free-threshing tetraploids, wild emmer, domesticated emmer to landrace across the genome. The landrace was divided into EA-, WA- and EU-landraces based on geographic distribution. Yellow lines represent introgression to EA-landrace, pink lines represent introgression to WA-landrace, and green lines represent introgression to EU-landrace. Grey dots are the chromosome centromeres.

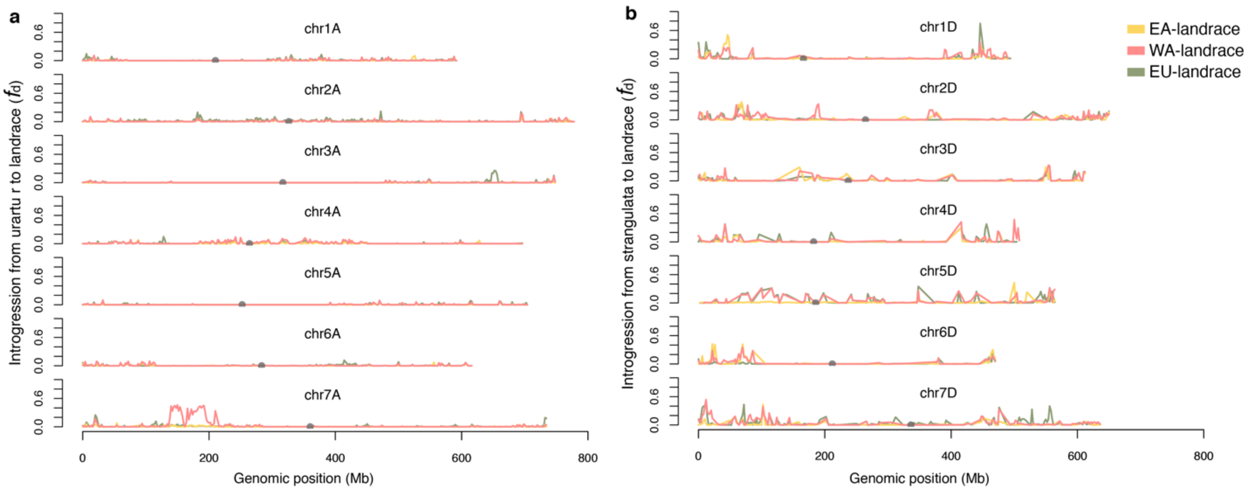


**Supplementary Fig. 10.** Introgression from diploid ancestors to landrace across the genome. **a,** The introgression from urartu to landrace. **b,** The introgression from Strangulata to landrace. The landrace was divided into EA-, WA- and EU-landrace based on geographic distribution. Yellow lines represent introgression to EA-landrace, pink lines represent introgression to WA-landrace, and green lines represent introgression to EU-landrace. Grey dots are the chromosome centromeres.

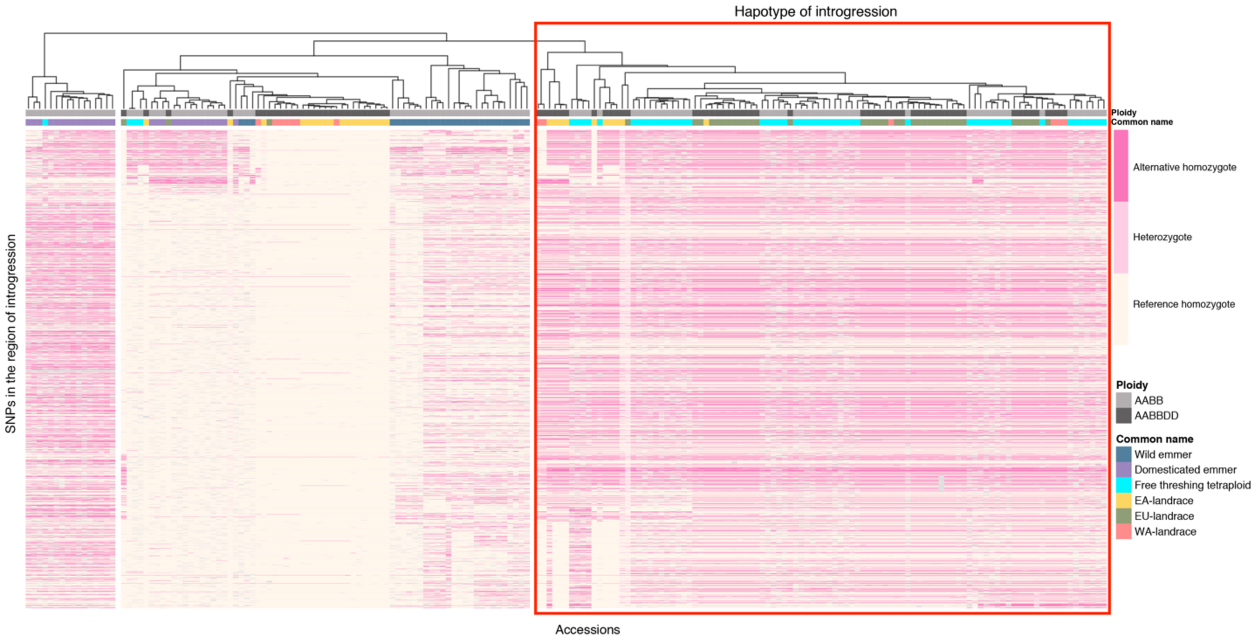


**Supplementary Fig. 11.** Haplotype of specific introgression region on Chr2A. The haplotype of introgression is showed in the red rectangle. The haplotype on Chr2A from position 165Mb to 540Mb was shown for each accession. Each row represents a genomic position for all accessions, and the column represents the haplotype of each accession. The ploidy of each accession is presented at the bottom of the cluster tree with different colors. The gray color represents the tetraploid wheat and the black color represents the hexaploid wheat. The different taxa are represented by different colors at the bottom of ploidy. Dark pink represents alternative homozygous, light pink represents heterozygous and light yellow represents reference homozygous.

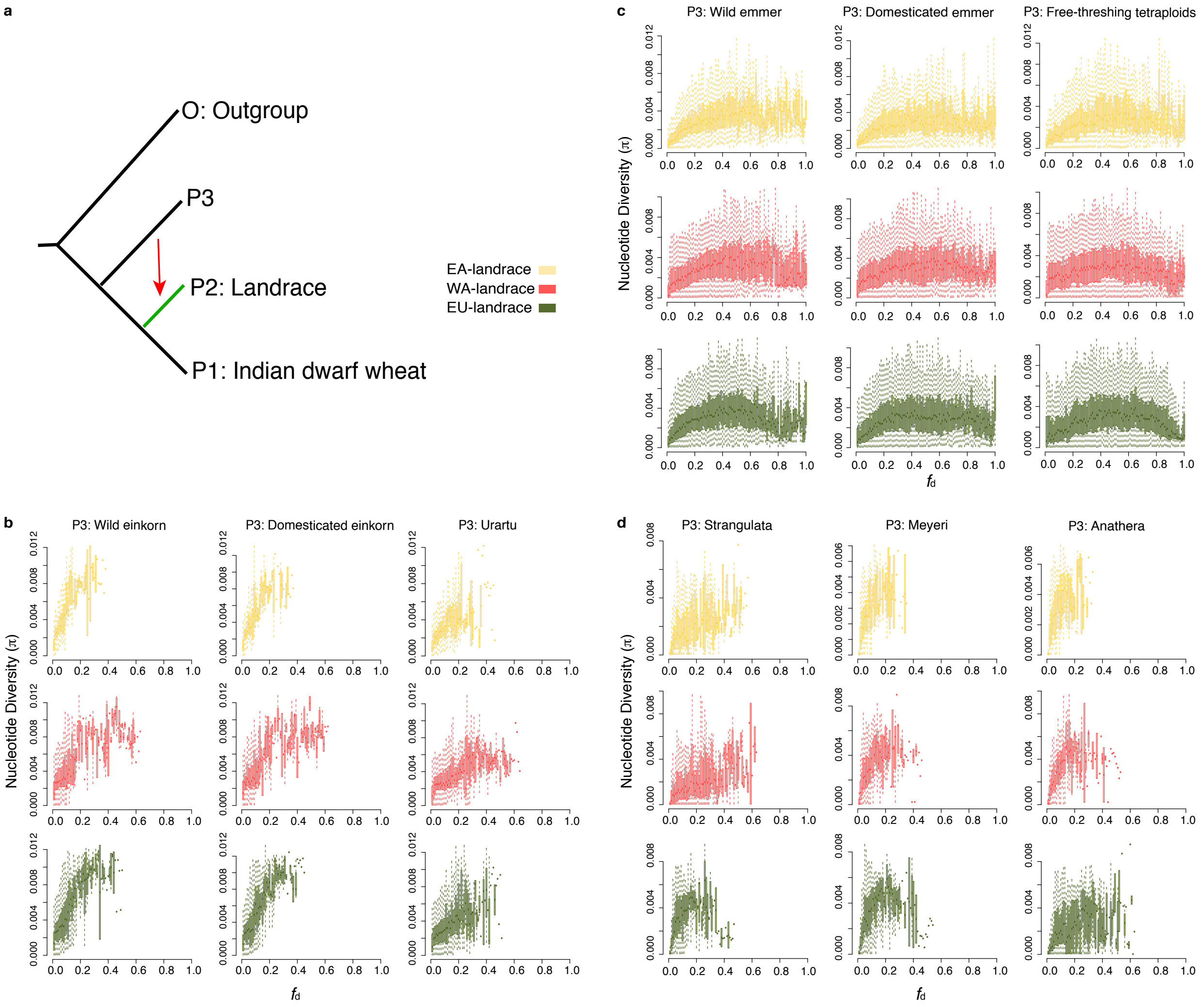


**Supplementary Fig. 12.** Nucleotide diversity is a function of $f_d$. **a,** The topology used for calculating $f_d$. **b,** Diploid species/subspecies in A lineage were used as P3. **c,** Tetraploid wheats were used as P3. **d,** Diploid species/subspecies in D lineage were used as P3.

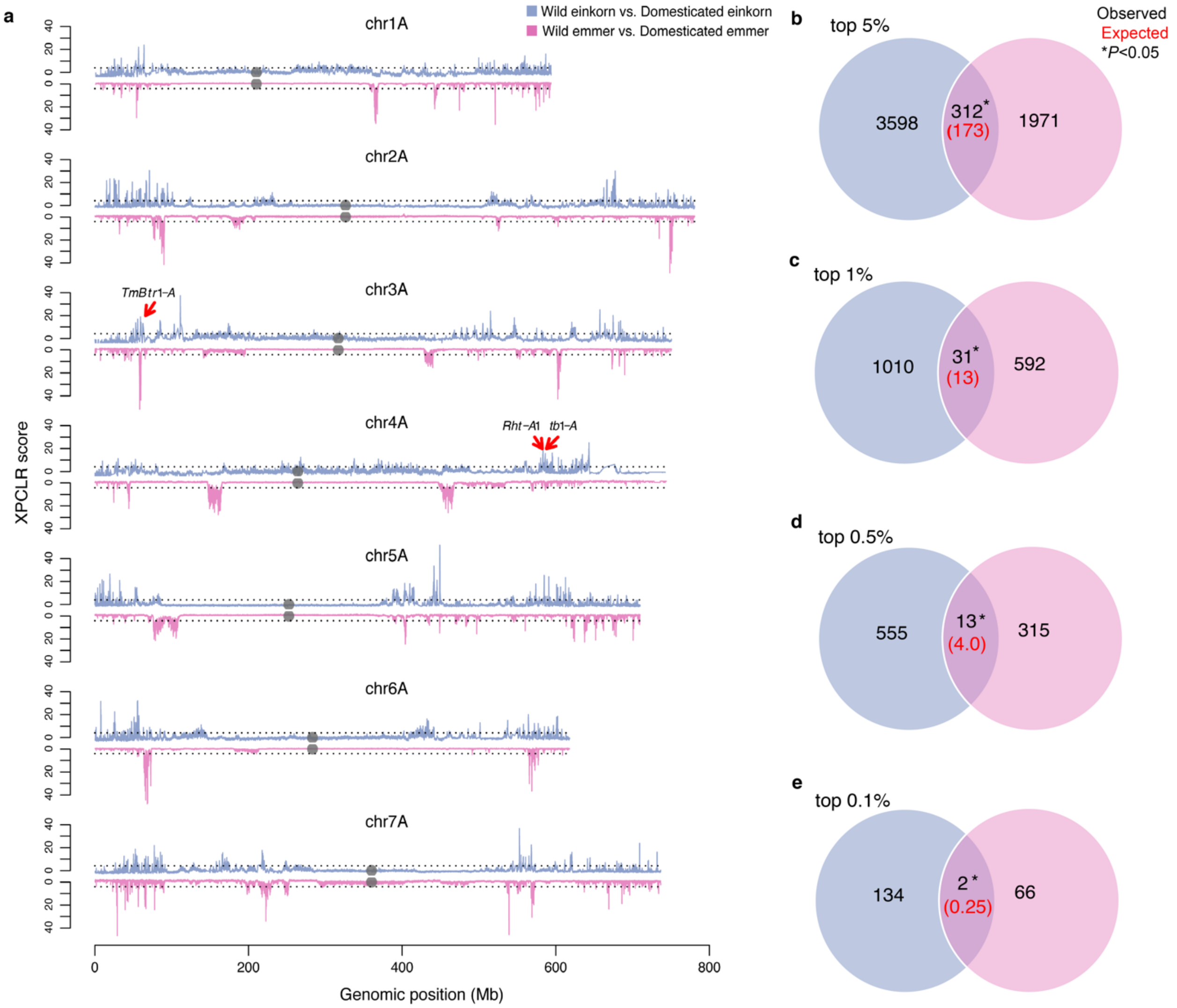


**Supplementary Fig. 13.** Comparison of selection sweeps detected from the paired domestication events. **a,** Selection sweep distribution across the genome. **b-e,** Convergence of selection with different threshold of XP-CLR score from top 5%, top 1%, top 0.5%, and top 0.1%. Blue lines or circles represent the XP-CLR score of domestication from wild einkorn to domesticated einkorn. Pink lines or circles represent the XP-CLR score of domestication from wild emmer to domesticated emmer. Gray dots represent the positions of centromeres. Number in red brackets represent the theoretical number under permutation test.

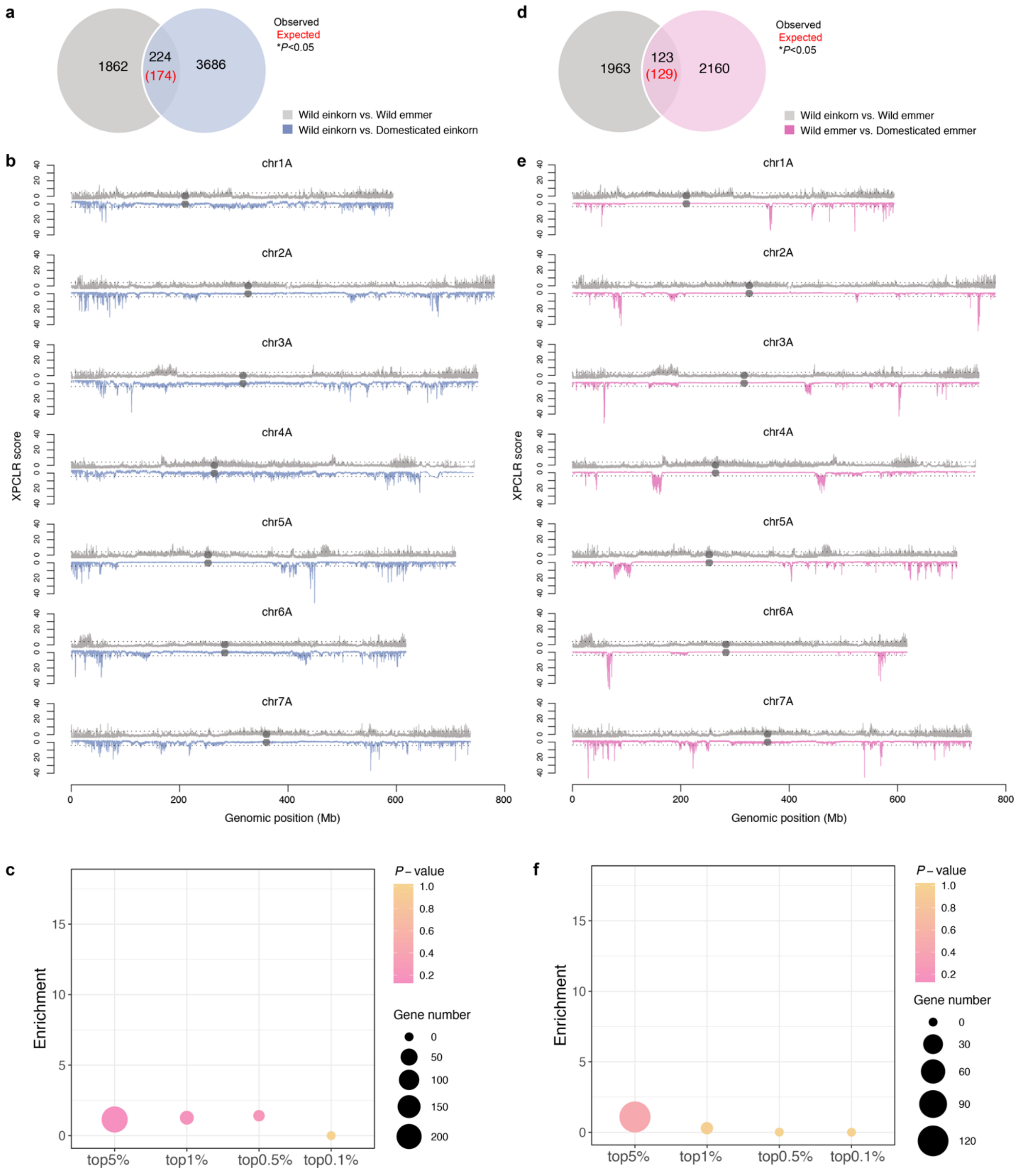


**Supplementary Fig. 14.** Selection sweeps comparison between negative control and domestication pairs. **a-c,** Comparison of genes selected between negative control (from wild einkorn to wild emmer) and domestication process from wild einkorn to domesticated einkorn. **d-f,** Comparison of genes selected between negative control and domestication process from wild emmer to domesticated emmer. (a) and (d) show the number of genes overlapped under the threshold of top 5%, (b) and (e) shows the whole genome distribution and (c) and (f) show the enrichment of convergence.

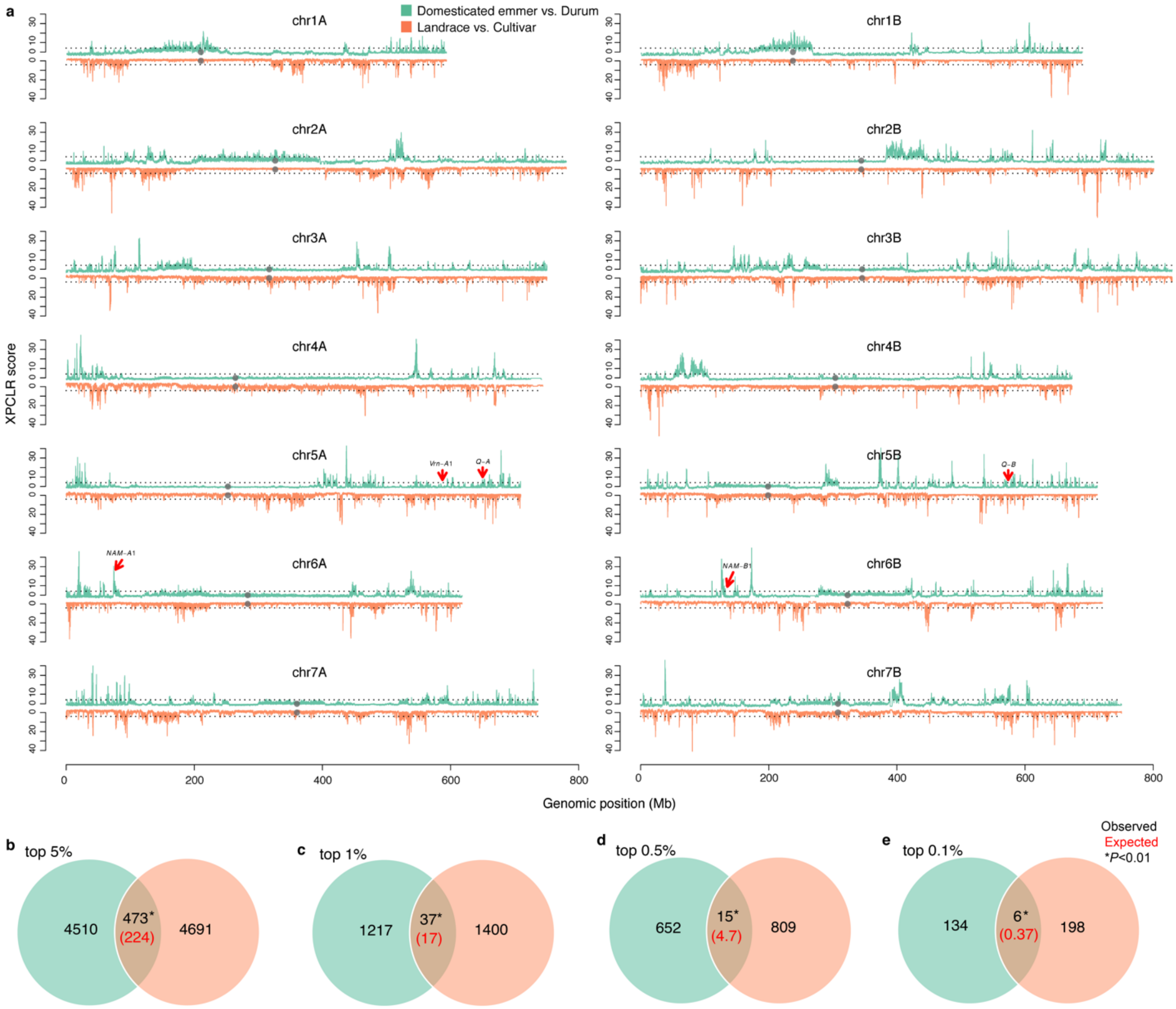


**Supplementary Fig. 15** Comparison of selection sweeps detected from the paired improvement events. **a,** Selection sweep distribution across the genome. **b-e,** Convergence of selection with different threshold of top 5%, top 1%, top 0.5%, and top 0.1%. Green lines or circles represent the XP-CLR score of domestication from domesticated emmer to durum. Red lines or circles represent the XP-CLR score of domestication from landrace to cultivar. Gray dots represent the positions of centromeres. Number in red brackets represent the theoretical number under random conditions.

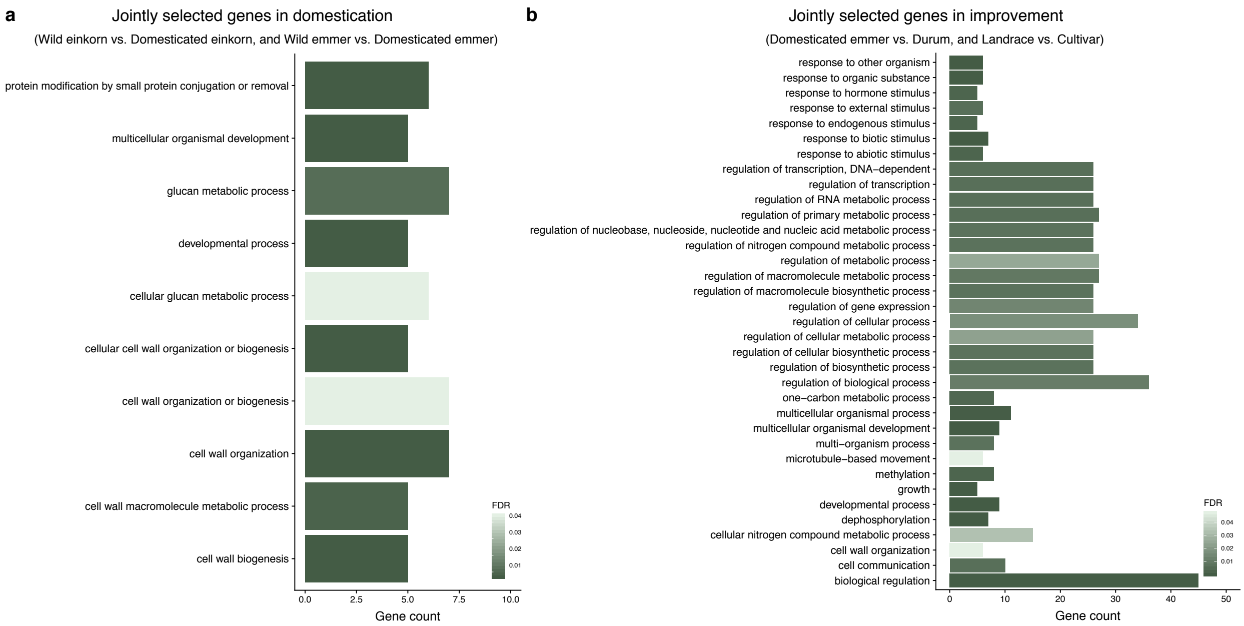


**Supplementary Fig. 16** GO enrichment for jointly selected genes. **a,** Enrichment for jointly selected genes in paired domestication events. **b,** Enrichment for jointly selected genes in paired improvement events. Only processes with five or more genes enriched were plotted.
